# Changes in the structure of spontaneous speech predict the disruption of hierarchical brain organization in first-episode psychosis

**DOI:** 10.1101/2023.12.12.570989

**Authors:** Rui He, Maria Francisca Alonso-Sánchez, Jorge Sepulcre, Lena Palaniyappan, Wolfram Hinzen

## Abstract

Psychosis implicates changes across a broad range of cognitive functions. These functions are cortically organized in the form of a hierarchy ranging from primary sensorimotor (unimodal) to higher-order association cortices, which involve functions such as language (transmodal). Language has long been documented as undergoing structural changes in psychosis. We hypothesized that these changes as revealed in spontaneous speech patterns may act as readouts of alterations in the configuration of this unimodal-to-transmodal axis of cortical organization in psychosis. Results from 29 patients with first-episodic psychosis (FEP) and 29 controls scanned with 7T resting-state fMRI confirmed a compression of the cortical hierarchy in FEP, which affected metrics of the hierarchical distance between the sensorimotor and default mode networks, and of the hierarchical organization within the semantic network. These organizational changes were predicted by graphs representing semantic and syntactic associations between meaningful units in speech produced during picture descriptions. These findings unite psychosis, language, and the cortical hierarchy in a single conceptual scheme, which helps to situate language within the neurocognition of psychosis and opens the clinical prospect for mental dysfunction to become computationally measurable in spontaneous speech.

## Introduction

Cognitive impairment is a well-established feature of schizophrenia, which is seen as early as premorbid stages and first-episode psychosis (FEP) and persists in the chronic phase of the illness (1). Although deficits in cognitive domains that employ linguistic representations such as verbal memory and processing speed (e.g., digital symbol coding ability) are most pronounced (2), no single cognitive domain appears to be fully spared in this illness, suggesting changes in the entire cognitive architecture from lower to higher-order functions (3, 4). Bottom-level sensorimotor disturbances, affecting a range of primary cortical functions such as audition and vision (5–8), may propagate upstream when integrative higher-order functions are employed (9). The implementational architecture for such integration has recently been studied with a nonlinear decomposition technique, diffusion mapping, which represents the very high-dimensional functional connectivity (FC) patterns across the cortex through a small number of continuous low-dimensional ‘gradients’ (10). Such gradients are considered to represent functional continuity and its variability across fundamental axes of organization of the human brain (11).

The first or principal gradient (G1) captures a cortical hierarchy ranging from functionally specialized sensorimotor cortex to domain-general association cortex, specifically including regions of the default mode network (DMN). Along this hierarchy, information is processed in increasingly abstract multimodal, heteromodal, and transmodal representational formats, forming a substrate for integrated higher-order cognition including language (12). The second gradient (G2) captures the functional separation of different primary sensory cortices, with visual cortex at one end and somatosensory cortex at the other. Gradients as proxies of hierarchical information streams can be further characterized with stepwise functional connectivity (SFC), a graph-based method to detect both direct and indirect functional links emanating from specific regions in the brain. Cortex-wise SFC, starting from sensory sources and culminating in the cortical hub of default mode regions (13), has already offered insight into the ‘bottom-up’ dysregulation of the hierarchy in schizophrenia, complementing gradient analysis (9).

The DMN is conceived as a dynamic ‘sense-making’ network, in which extrinsic incoming information is integrated with prior intrinsic information, so as to construct rich situation models with narrative meaning, which unfolds over time (14). This conception suggests that the particular hierarchy captured by G1 may functionally relate to meaning-making as well, i.e., the building of higher-order meaning from perception, in which language plays a role. We hypothesized that a disrupted topological architecture of the cortex in psychosis (9) would be reflected in structural language changes, which have also long been observed in psychosis (15, 16).

Specifically, speech production necessarily involves retrieving concepts internally from semantic memory. Out of context, such concepts associate with each other at certain semantic distances, with *cat* being relatively more similar semantically to *kitten* than to *house*. These distances can be computationally quantified through cosine similarities between word embeddings retrieved from large neural language models (LMs) (17). The meaning of language, however, depends on relating words grammatically, whereby they become parts of phrases (e.g., a noun phrase), and these phrases become parts of other phrases (e.g., a verb phrase), giving rise to a syntactic hierarchy. At this level, language can encode full interpretations of perceptual experience, as in the episodic thought *It was the cat*, which will typically encode reference to a specific event in space and time and form part of a narrative. Both lexical-semantic associations and syntax can be studied through graphs, using advanced natural processing tools. We reasoned that a disruption in cortical gradients could be reflected in the graph-theoretic structure of meaning in language, during a simple speech task such as describing a picture. While numerous studies have used semantic distances between words (18–20) as well as syntactic complexity (21, 22) to quantify language changes in psychosis, relations between these and neurofunctional changes have barely been explored (but see (23, 24)).

We predicted alterations in meaning structure should bear on large-scale cortical gradients and SFC in psychosis. This is because language not only plays a crucial role in the creation of meaning but is also inherently integrative of cognition as whole: any utterance we make requires integration with perception, fine motor control, semantic memory, attention, and social cognition. Empirical support for this expectation comes from the fact that inter-individual differences in the geodesic distance of sensorimotor landmarks to the language and default mode networks covary with activations in these networks during language tasks (25). Furthermore, greater separation between the DMN and sensorimotor network is related to semantic cognition, especially our ability to retrieve links between concepts from semantic memory (26). In this context, the semantic brain network, which includes essential left-hemispheric default mode and limbic regions (27), is of particular interest, given its reported associations with language disruptions in psychosis (28, 29). The results of our study provide, for the first time, speech graph alterations related to aberrations in the large-scale cortical gradients with specific involvement of the semantic network in psychosis. The graphs reveal a semantic space altered both in terms of its lexical-conceptual semantic associations between individual units of meaning and of the syntactic structures that enable meaning at the sentential and narrative level. This finding contributes to our understanding of the functional significance of neurocognitive changes in psychosis and highlights spontaneous speech as a readout of these changes and a potential source for their clinical measurement.

## Results

### Altered cortical gradients and SFC

Figure 1A depicts the workflow of the fMRI analysis. Twenty-nine untreated first-episode psychosis (FEP) patients, matched to twenty-nine healthy controls (HC) on age, gender, and education, participated in this study (see Table 1). Subjects underwent resting-state fMRI scans on a Siemens 7T Plus (Erlangen, Germany) and were asked to describe three pictures from the Thematic Apperception Test (one minute for each picture) immediately prior to the scans. FC matrices were extracted from 1000 regions-of-interest ROIs using the Schaefer 1000-parcellation (30) for each subject and group. Affinity matrices were computed from these, where affinity values between any pair of regions are the cosine similarities between the vectors representing these regions in terms of their FC to all other cortical regions. Diffusion embedding was then applied to these affinity matrices to extract ten group-level gradients from these affinity matrices, resulting in ten gradient values (i.e., affinities) assigned to each parcel (31). Group templates for all gradients with scree plots showing the variance explained by each gradient, as well as those for SFC analyses, are available in Fig. S12-S24. Subject-level gradient templates were aligned to their corresponding group templates. Gradient values of ROIs were compared between HC and FEP using surface-based linear models (SLM).

**Figure 1.**
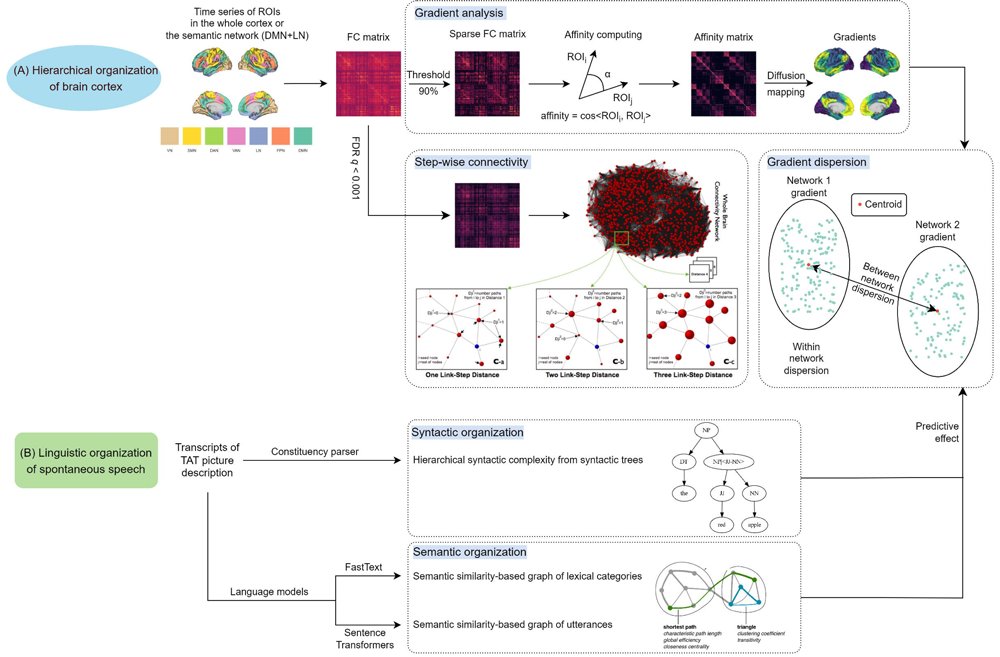
Workflow of this study. To determine the cortical hierarchy, we carried out gradient analysis in the whole cortex and within the semantic subnetwork, with gradient dispersion derived from it, and stepwise functional connectivity (SFC) to better interpret gradient results. The illustration graph for SFC is adapted from Sepulcre et al. (13) For linguistic organization of spontaneous speech, we computed formal syntactic measures and lexical-conceptual semantic graph-theoretical measures, the latter of which capture shortest paths, clustering coefficients, and the balance between them (small world organization). The visualization of these graph-theoretical measures is adapted from Rubinov and Sporns (65).

**Table 1.** Descriptive statistics of the dataset.

|  | HC | FEP | Test | *p* | Effect size |
| --- | --- | --- | --- | --- | --- |
| Num | 29 | 29 | / | / |  |
| Age | 21.5 (3.6) | 21.4 (2.2) | Welch's *T* test | 0.861 | -0.046 |
| Sex | 24.1% | 27.6% | $\chi^2$ test | 0.764 | 0.039 |
| Education Level | 71.4% | 51.7% | $\chi^2$ test | 0.127 | 0.198 |
| Parental SES | 58.0% | 45.0% | $\chi^2$ test | 0.139 | 0.196 |
| P1: Delusions | / | 5 (2) | / | / | / |
| P2: Conceptual disorganization | / | 3 (3) | / | / | / |
| P3: Hallucinatory behavior | / | 5 (1) | / | / | / |
| N1: Blunted affect | / | 2 (3) | / | / | / |
| N4: Passive/apathetic social withdrawal | / | 3 (4) | / | / | / |
| N6: Lack of spontaneity and flow of conversation | / | 1 (2) | / | / | / |
| G5: Mannerisms & posturing | / | 1 (2) | / | / | / |
| G9: Unusual thought content | / | 4 (2) | / | / | / |
| Total TLI socre | 0.30 (0.30) | 1.51 (1.30) | Welch's *T* test | < .001 | -1.226 |

Note: Age was indicated by mean ± standard deviation with effect size indicated by Cohen's *d*. Sex: proportion of female subjects. Education level: proportion of subjects receiving formal education of over 12 years. Parental social economic state (SES): proportion of subjects with parental SES less than 3. TLI: thought and language index. Effect sizes for group comparison on sex and education were indicated by contingency coefficient. Clinical scores for FEP were indicated by median (IQR, interquartile range). Statistical tests within this table were carried out with JASP 0.16.4.

Figure 2 summarizes the gradient findings. Figure 2A shows an overall (global) gradient compression in FEP, with a smaller range and a higher peak, suggesting a decreased functional distance between the endpoints of cortical hierarchies. More specifically, when plotting gradient values of G1 and G2 together, both groups show an expected geometry (see Figure 2B), with sensorimotor networks maximally separated from the DMN in the first gradient (G1, vertical axis), and the visual network (VN) and somatomotor network (SMN) maximally separated from each other in the second gradient (G2, horizontal axis). However, in G1, the functional distance between the DMN on top and the VN at the bottom *decreases* in FEP, while that of the DMN and the SMN *increases*. Moreover, in G2, the dispersion between VN and SMN is maintained, while, between them, the DMN is less distant from the SMN and more distant from the VN. Group templates of all ten cortical gradients can be found in Fig. S12-S15. In line with this pattern, Figure 2C shows that for G1 the gradient values of ROIs in VN significantly increased in FEP while those in SMN significantly decreased. For G2, gradient values of ROIs in the two extremes, VN and SMN, were unchanged, while gradient values in DMN significantly decreased in FEP.

**Figure 2.**
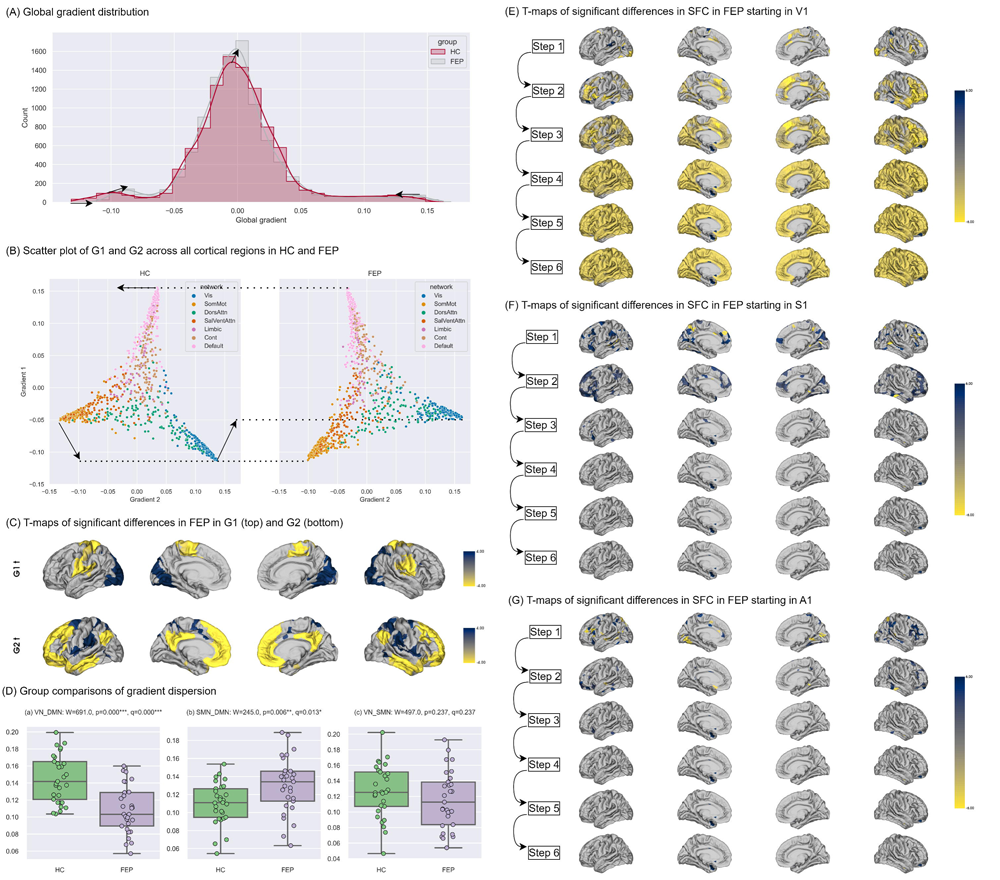
Summary of changes in cortical functional hierarchy. (A) Histogram with Global distribution of all ten cortical gradients, with a density line estimated by kernel density estimator (KDE) and arrows showing changes in both the peaks and bilateral ends of the density lines in first-episode psychosis (FEP) relative to health controls (HC). Red: HC; grey: FEP. (B) Scatter plots of the first (y-axis) and the second (x-axis) gradient in HC (left) and FEP (right). The color coding refers to Yeo’s 7 networks: blue: visual (Vis); orange: somatomotor (SomMot); green: dorsal attention (DosAttn); red: salience/ventral attention (SalVentAttn); purple: limbic (Limbic); brown: frontoparietal control (Cont); pink: default mode (Default). Arrows indicate how the three vertices of the triangular figure (visual, somatomotor, and default mode networks) change in FEP. (C) *T* values from the surface-based linear model (SLM) model plotted on the cortical surface, showing regions in FEP that differ from HC on the first (top) and second (bottom) gradients. Only *T* values with statistical significance (*q* < 0.05) are plotted. Blue indicates higher gradient values in FEP while yellow indicates lower values. The denser the color is, the larger the *T* values. (D) Group comparisons of gradient dispersion between (a) VN and DMN, (b) SMN and DMN, and (c) VN and SMN. Significance levels: *** 0.001, ** 0.01, * 0.05. (E-G) *T* values out from the SLM model showing the effect of being FEP on the seed-based step-wise functional connectivity (SFC) degrees at each of the six steps, for (E) primary visual seed (V1), (F) primary somatosensory seed (S1), and (G) primary auditory seed (A1). Only T values with statistical significance (*q* < 0.05) are plotted. Blue indicates higher gradient values in FEP while yellow indicates lower values. The denser the color is, the larger the T values.

For further confirmation of this pattern, we directly quantified the relative distance between networks using refined gradient dispersion from Bethlehem et al. (32). Dispersion between VN and DMN, as well as dispersion between SMN and DMN, was defined as the difference between the averaged first gradient values across ROIs within each network. Dispersion between SMN and VN was defined as the difference between the averaged second gradient values across ROIs within each network. As shown in Figure 2D, there was significantly lower VN-DMN dispersion in FEP (suggesting lesser functional separation between VN and DMN), while dispersion between SMN and DMN was higher in FEP (suggesting decreased affinity).

Seed-based SFC maps were calculated following Sepulcre et al. (13), using bilateral primary visual (V1), auditory (A1) and somatosensory (S1) ROIs as seeds. These maps show the evolution of counts of functional links between a ROI and the remainder of cortical regions as the stepwise distance (number of links) between them is increased. Group-level cortical SFC maps are available in Fig. S20-S22. Figures 2E-G show significant increases (blue) and decreases (yellow), in FEP relative to controls, as link-steps increase step by step (13). Specifically, in FEP, there were less links in FEP propagating from V1 to DMN regions and remaining regions of association cortex (Figure 2E), while, to a lesser extent, there were more links between S1 and DMN (Figure 2F), as well as between A1 and DMN (Figure 2G). These results confirm at the level of SFC, an alteration, more pronounced in the case of the visual seed, in the emergence of (or progression to) a trans-modal architecture relevant for higher-order functions such as language. Overall, the observation is that of decreased stepwise links between V1 and DMN along with a reduced dispersion between them, and an increase in stepwise connections between the somatosensory cortex and the DMN, in FEP.

### Compressed gradient within the semantic network

The same analysis pipeline as above was therefore applied within the semantic network, defined as comprising the left default and limbic regions following Binder et al. (27) specifically: (1) the frontal-parietal heteromodal association cortex to process heteromodal and multimodal information, mainly comprising the dorsomedial and dorsolateral prefrontal cortex (dmPFC, dlPFC), middle frontal gyrus (MFG), inferior frontal gyrus (IFG), the angular gyrus (AG), the adjacent supramarginal gyrus (SMG), and the lateral temporal lobe; and (2) the self-related medial limbic cortex, mainly comprising the ventromedial temporal lobe, ventromedial and orbital prefrontal cortex, and the posterior cingulate gyrus (PCC) and adjacent ventral precuneus (Prec)

(27). Group templates of all ten semantic gradients can be found in Fig. S16-S19 and Tables S3-S4. As shown in Figure 3A, we observed a principal semantic gradient ranging from lateral heteromodal association cortex (IFG, dlPFC, SMG/AG, and posterior middle temporal gyrus (pMTG)) (as seen in bright colors), to medial limbic regions (PCC, Prec) (as seen in dark colors). In FEP, this hierarchy was generally retained. However, as seen in Figure 3B-C, the high-end points of this hierarchy in the anterior PCC and medial temporal lobe lowered in FEP, while the low-end points (AG, SMG, pMTG, and dlPFC) ascended. This, together with a significantly lower within-network dispersion in FEP (Figure 3E), suggests a compressed semantic network, in which the overall hierarchy is reduced in FEP. Figure 3D further illustrates that the distribution of this gradient also became more centralized in FEP, changing from a left-skewed trimodal distribution to a bell-shaped unimodal distribution.

**Figure 3.**
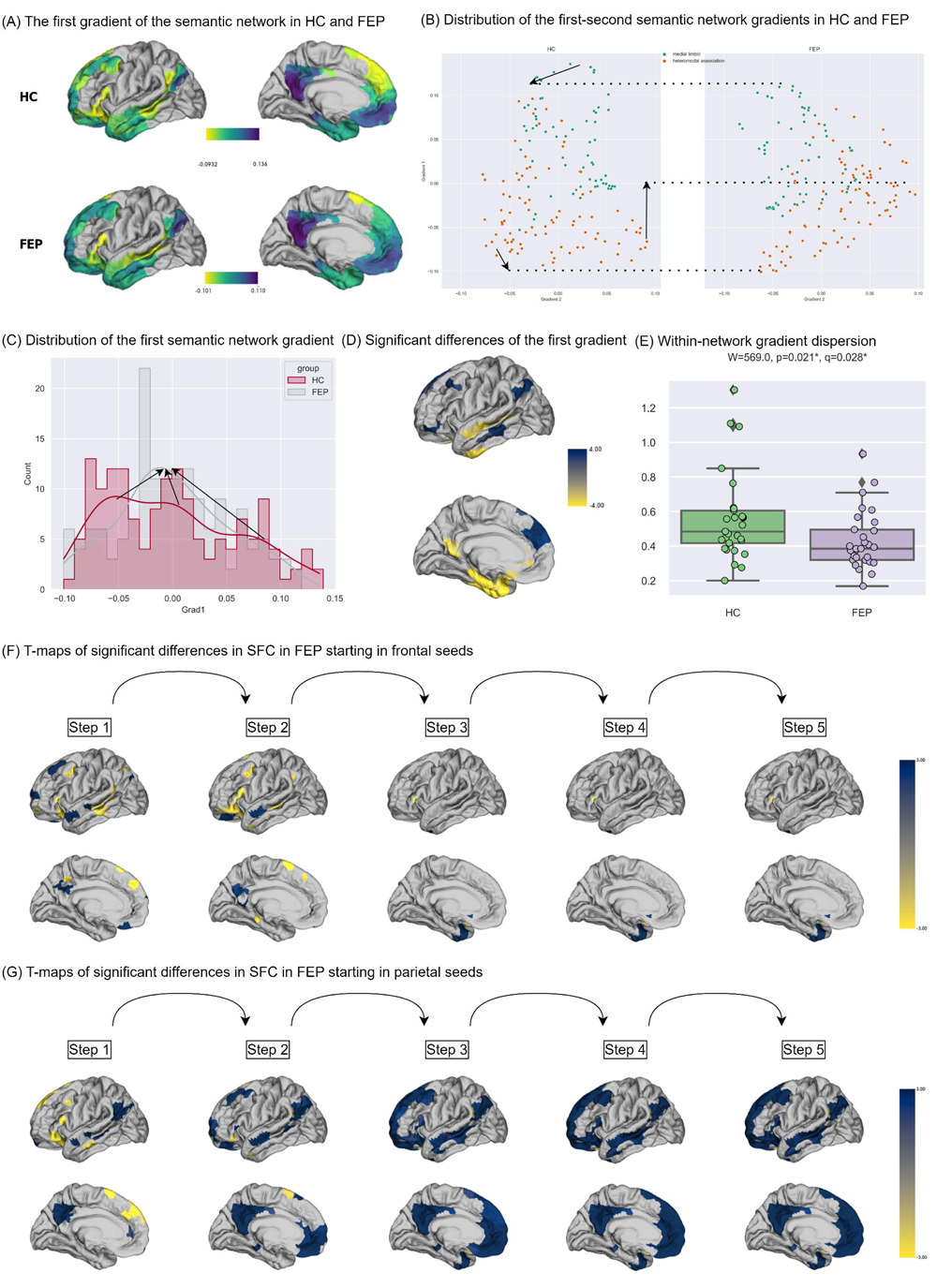
Summary of changes in the functional hierarchy within the semantic network. (A) Spatial map of the first gradient of the semantic network on cortical surface. (B) Scatter plots of the first (y-axis) and the third (x-axis) gradient in health controls (HC, left) and first-episode psychosis (FEP, right). The color refers to 2 cortices in the semantic network as described in the main text: orange: heteromodal association cortex; green: medial limbic cortex. (C) Histogram with global distribution of the first semantic network gradient with density line and arrows showing changes in the peaks and bilateral ends of the density line from HC to FEP. Red: HC; grey: FEP. (D) *T* values out from the surface-based linear model (SLM) model showing the effect of being FEP on the first gradient of the semantic network. Only *T* values with statistical significance (*q* < 0.05) are plotted. Blue indicates higher gradient values in FEP while yellow indicates lower values. The denser the color is, the larger the *T* values. (E) Group comparisons of the within-network dispersion of the first semantic network gradient. (F,G) *T* values out from the SLM model showing the effect of being FEP on the seed-based step-wise functional connectivity (SFC) degrees at each of the five steps, for (F) seeds for the frontal heteromodal association cortex (IFG and pMTG) and (G) seeds for the parietal heteromodal association cortex (AG and SMG). Only *T* values with statistical significance (*q* < 0.05) are plotted. Blue indicates higher gradient values in FEP while yellow indicates lower values. The denser the color is, the larger the *T* values.

Again, we followed up on this gradient analysis with a SFC analysis, choosing as seeds four ROIs at the ‘low’ end of the hierarchy as depicted in Figure 3B: the frontal seeds IFG and dlPFC, on the one hand, and the parietal ones, supramarginal gyrus (SMG) and angular gyrus (AG), on the other. Group-level SFC maps within the semantic network are available in Fig. S23-S24. Figure 3F-G illustrates increased links between the two ends of the principal gradient in FEP. Specifically, SMG and AG had more links to the whole semantic network in FEP. Moreover, the IFG and dlPFC had more connections to medial limbic regions but fewer links to these two ROIs themselves.

### Behavioral linguistic results

Several findings of changes in semantic cognition in psychosis (16, 18–20, 33–35) motivate the prediction of a relation between semantic structure changes in spontaneous speech and the changes in gradients observed above. We constructed semantic graphs to represent the semantic structure of discourse produced during picture descriptions at two levels, based on the associations among units of meaning, which were either content words (*woman*, *farm*, *horse*, etc.) or utterances. Word meanings were represented through embeddings derived from the FastText (FT) model pre-trained on English data (36) (cc.en.300.bin). We computed binarized sparsified semantic similarity graphs using the normalized cosine similarity scores between every pair of two embeddings, from which graph-theoretical measures were extracted showing centrality, clustering, and the balance between these two in the small-worldness coefficient, as specified in Table 2. Following a similar procedure, we embedded every utterance using a sentence-transformers (ST) model (37) (https://huggingface.co/sentence-transformers/all-MiniLM-L6-v2), and constructed similar semantic graphs with utterances as the nodes. As shown in Figure 4B, despite a similar quantity of lexical categories in both groups, both semantic graphs (of content words and of utterances), became more ‘concentrated’ in FEP, as indicated by higher closeness centrality and global efficiency. The balance between graph-wise segregation of semantic units (centrality) and integration of closely related units (clustering), as indicated by the small-worldness coefficient (sigma, *σ*), was significantly lower in FEP.

**Figure 4.**
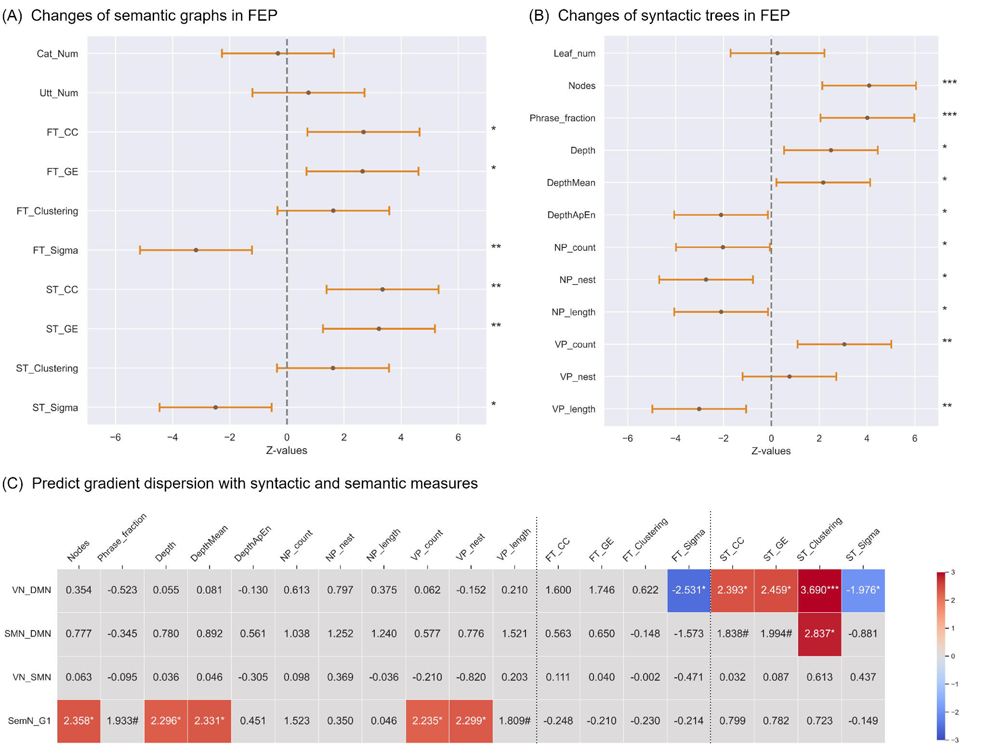
Linguistic changes and their relations to gradient dispersion. Significance levels: *** 0.001, ** 0.01, * 0.05. (A, B) Line plots indicating the semantic and syntactic changes in FEP, respectively. Asterisks on the right denotes the levels of statistical significance. Each error bar featured a dot representing the *z* value, while the length of the bar represented the 95% confidence interval. The dash lines anchored *z* values of zero. Error bars with the dot falling left to the line signified an increase in FEP while falling right to the line signified a decrease in FEP. (C) Heatmaps indicating the predictive effect of linguistic measures on gradient dispersion between VN and DMN, SMN and DMN, VN and SMN, and the intrinsic dispersion of semantic network (SemN_G1) via the dispersion of its first gradient. Red cells signified positive predictive effects while blue cells signified negative predictive effects. Only cells where significant predictive effects were observed (corrected for False Discovery Rate with *q* < 0.05) were colored. Moreover, we also marked those cells with *q* in the interval of [0.5, 0.1) with the number sign (#). FT for FastText-based measures and ST for Sentence transformers. NP: noun phrases; VP: verb phrases; ApEn: approximate entropy; VN: visual network; SMN: somatomotor network; DMN: default mode network; SemN: semantic network; GE: global efficiency; CC: closeness centrality.

**Table 2.**
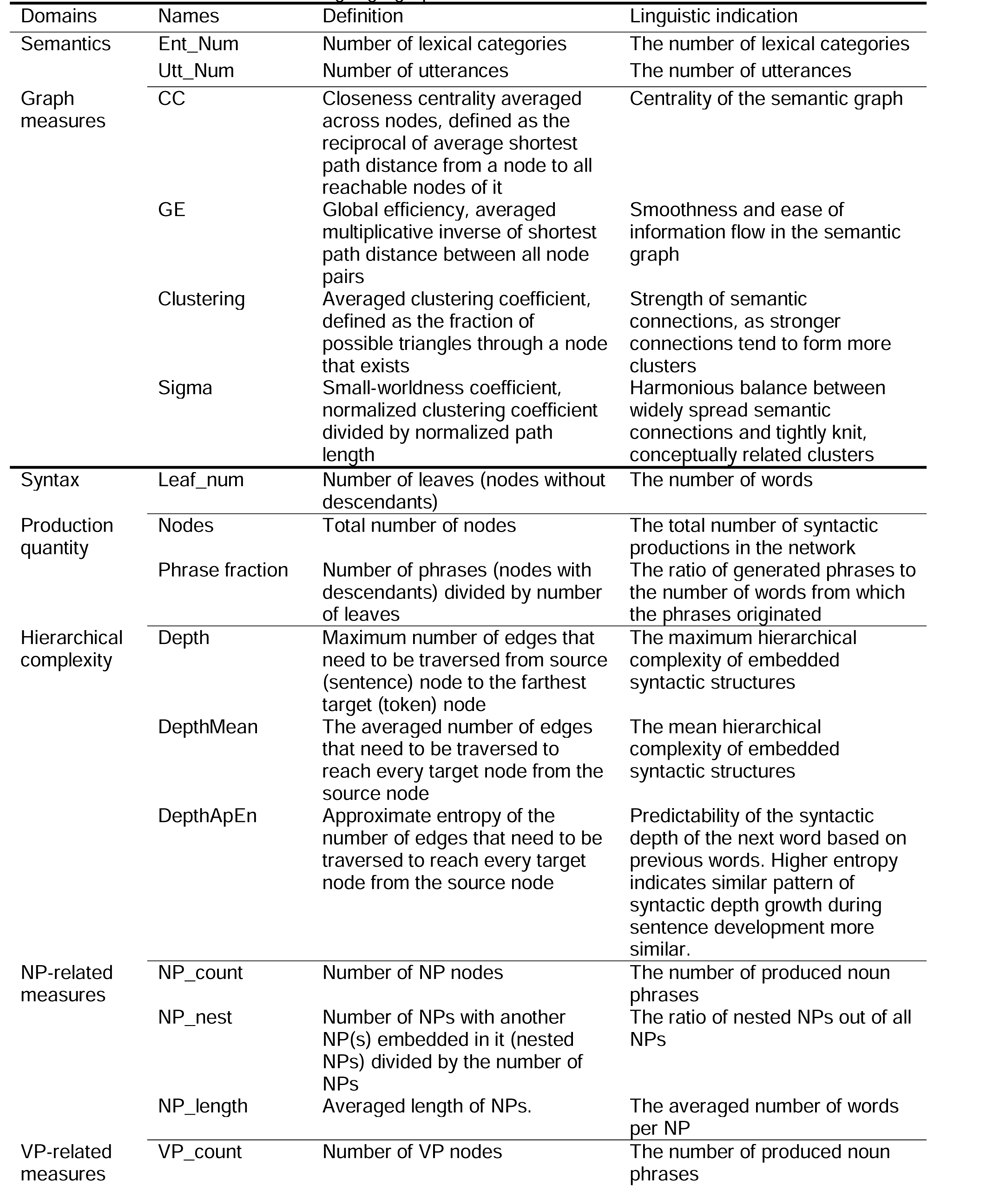
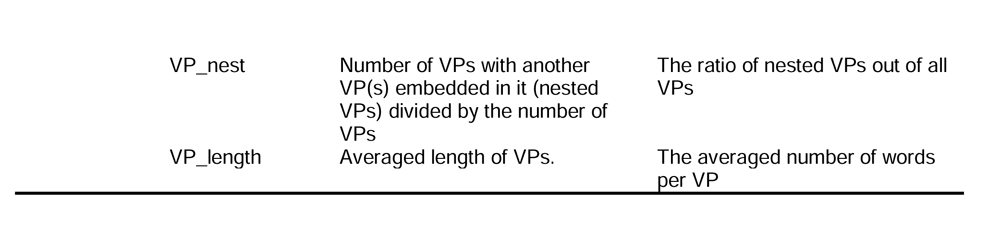
Features extracted from language graphs.

| Domains | Names | Definition | Linguistic indication |
| --- | --- | --- | --- |
| Semantics | Ent_Num | Number of lexical categories | The number of lexical categories |
|  | Utt_Num | Number of utterances | The number of utterances |
| Graph measures | CC | Closeness centrality averaged across nodes, defined as the reciprocal of average shortest path distance from a node to all reachable nodes of it | Centrality of the semantic graph |
|  | GE | Global efficiency, averaged multiplicative inverse of shortest path distance between all node pairs | Smoothness and ease of information flow in the semantic graph |
|  | Clustering | Averaged clustering coefficient, defined as the fraction of possible triangles through a node that exists | Strength of semantic connections, as stronger connections tend to form more clusters |
|  | Sigma | Small-worldness coefficient, normalized clustering coefficient divided by normalized path length | Harmonious balance between widely spread semantic connections and tightly knit, conceptually related clusters |
| Syntax | Leaf_num | Number of leaves (nodes without descendants) | The number of words |
| Production quantity | Nodes | Total number of nodes | The total number of syntactic productions in the network |
|  | Phrase fraction | Number of phrases (nodes with descendants) divided by number of leaves | The ratio of generated phrases to the number of words from which the phrases originated |
| Hierarchical complexity | Depth | Maximum number of edges that need to be traversed from source (sentence) node to the farthest target (token) node | The maximum hierarchical complexity of embedded syntactic structures |
|  | DepthMean | The averaged number of edges that need to be traversed to reach every target node from the source node | The mean hierarchical complexity of embedded syntactic structures |
|  | DepthApEn | Approximate entropy of the number of edges that need to be traversed to reach every target node from the source node | Predictability of the syntactic depth of the next word based on previous words. Higher entropy indicates similar pattern of syntactic depth growth during sentence development more similar. |
| NP-related measures | NP_count | Number of NP nodes | The number of produced noun phrases |
|  | NP_nest | Number of NPs with another NP(s) embedded in it (nested NPs) divided by the number of NPs | The ratio of nested NPs out of all NPs |
|  | NP_length | Averaged length of NPs. | The averaged number of words per NP |
| VP-related measures | VP_count | Number of VP nodes | The number of produced noun phrases |

|  |  |  |
| --- | --- | --- |
| VP_nest | Number of VPs with another VP(s) embedded in it (nested VPs) divided by the number of VPs | The ratio of nested VPs out of all VPs |
| VP_length | Averaged length of VPs. | The averaged number of words per VP |

At the sentence level, meaning arises through syntactic structure. We therefore tested how lexical concepts were organized into meaningful sentences based on formal syntactic rules rather than their semantic associations with other lexical concepts (as in the above graphs). Syntactic structures geometrically take a hierarchical form typically represented as trees, within which words are organized into phrases (e.g., noun phrases, NPs), which become parts of other phrases (e.g., verb phrases, VPs), until the top node (the S node representing the sentence itself) is reached. The measures extracted from syntactic trees are specified in Table 2.

We first extracted the total number of nodes in the tree, as well as the ratio of generated phrases to the number of words from which the phrases originated, as indicators for production quantity. Both indicators were significantly higher in FEP than HCs. Then, we defined the syntactic depth of each word as the number of edges in the syntactic tree that need to be crossed from this word to the sentence node. From this we obtain a time series of syntactic depth values from the first to the last word of a sentence. Both the maximum of such arrays and the averaged word-level depth *increased* in FEP. The approximate entropy, which indicates the predictability of the next syntactic depth from the previous ones, however, *decreased* in FEP. The number and averaged length of NPs, as well as the ratio of nested NP, also significantly decreased in FEP. On the other hand, the number of VPs significantly increased, though with a significant decrease in their average length. In short, with more but shorter predicative structures produced (VPs), FEP filled the argument positions of these predicate structures with fewer, shorter, and simpler NPs.

### Linguistic metrics predict cortical hierarchies

Generalized linear models (GLM) were applied to predict gradient dispersion from semantic graph measures. As shown in Figure 4C, the dispersion between VN and DMN was negatively predicted by both FastText (FT)-and sentence transformer (ST)-based small-worldness coefficients (sigma, *σ*), and positively by ST-based graph measures of centrality, efficiency, and clustering. In turn, as seen in Figure 4C, the number of words and nodes, the maximum and mean syntactic depth, and VP measures significantly predicted the dispersion of the semantic brain subnetwork, even after correcting for multiple comparisons, in addition to two noteworthy tendencies (*q* < 0.1): phrase fraction (*z* = 1.933, *q* = 0.053) and averaged length of VP (*z* = 1.809, *q* = 0.070).

## Discussion

This study investigated changes in the hierarchical organization of cortical functions in FEP through gradients of functional connectivity and how these relate to the semantic organization of spontaneous speech at lexical and structural levels. Firstly, the results confirm a reorganization of cortical hierarchies in FEP. This includes both the first or principal gradient (G1), which represents the functional spectrum ranging from direct perception and action towards the integration and abstraction of information in a multi-to transmodal format, and the second gradient (G2), which separates two primary sensorimotor cortices at its two ends, namely the VN and SMN, respectively (10). Specifically, the functional distance in G1 between VN and the DMN was compressed in FEP, while that between SMN and DMN was decompressed. In G2, the DMN moved from its position in the middle of the neurotypical G2 hierarchy towards a greater functional distance from the VN and a lesser such distance to the SMN. Together, these large-scale shifts in network hierarchies depict a disturbance of the balance between sensorimotor and higher-order association cortices in schizophrenia, which were previously documented by Dong et al. (9). The shifts are consistent with the SFC analysis, which also demonstrates an uncommon clustering of sensorimotor cortices and the DMN in FEP in terms of the direct and indirect links between them. In particular, along with a decreased dispersion between the VN and the DMN in G1, stepwise connections between V1 and DMN decreased as well. In contrast, stepwise connections increased along with an increased dispersion between SMN and DMN in G1. Overall, the SFC pattern is that there is greater clustering of SMN and DMN and weaker clustering between VN and DMN as well as VN and SMN regions.

Our key question was how such tightening and loosening of the integration between hierarchically ordered cortices is reflected in language, viewed as a higher-order cognitive function that depends on the integration of multiple cognitive systems. The graphs of both word-level and utterance-level meaning (Figure 4B) demonstrate a pattern of significantly higher centrality, weaker small-worldness, and a tendency for greater clustering. We capture this conceptually as a ‘shrinking’ or ‘narrowing’ semantic space, consistent with recent findings, which found a lesser average semantic distance (higher cosine semantic similarity) between consecutive words in FEP (19, 38). Higher centrality in these graphs suggests excessive connections between semantically distant units, while higher clustering indicates the formation of more semantic subdomains. Put differently, the flow of meaning in the speech produced by FEP develops over more nuanced topic structures, with more semantic boundaries (of clusters) within a shrinking semantic space.

These semantic graph properties predicted the separation between primary sensory and higher-order cognition networks in the principal cortical gradient. Since this gradient reflects the dispersion between peripheral-perceptual and higher-order cortices, this relation may indeed reflect the specific task demand in a picture description: converting perceptual (visual) information into language, translating percepts into higher-order meaning as carried by language.

Related work already suggests that greater physical distances between DMN regions and sensorimotor cortices may be essential for cognitive processes drawing on semantic memory (25, 26). Going beyond these previous studies, the present one shows a relation between gradients and the semantic organization of spontaneous speech as assessed with neural language models. In turn, Smallwood et al. (39) developed a topographical perspective on the functionality of the DMN, noting that, compared to other unimodal areas, the primary auditory cortex is relatively proximal to key DMN and language processing regions such as the angular, inferior frontal, and middle temporal gyri. They speculate that ‘this proximity to the auditory system allows these regions of the DMN to capitalize on the capacity for language processes to organize cognitive function, perhaps through the vehicle of inner speech’. This intriguing suggestion is pertinent in the context of psychosis, where auditory verbal hallucinations, among other symptoms, reflect a disruption of a (ordinarily silent) process of inner speech. Increased links between the auditory and somatosensory seeds and DMN in our study are, in this respect, an intriguing lead to follow up on in future studies, which might target a group of voice hearers specifically.

In the semantic brain network, we identified a new principal gradient separating the multi-and heteromodal association cortex in lateral frontotemporal and parietal areas, from medial limbic regions, specifically the PCC and the parahippocampal gyrus. The hierarchy captured by this gradient thus ranges from the frontotemporal subdivision of the semantic network, functioning as information storage, retrieval, and update, to medial limbic regions, which are related to the encoding of episodic information including emotions and social interactions (27, 40). This latter information is critically related to the internal self and its narrative (27), potentially indicating that the semantic gradient relates to the process of integrating perceptual and conceptual semantic information from the external world to episodic memory and the self. Remarkably, this hierarchy of semantic processing, based on resting-state data, aligns with a recent semantic tiling of the cortex derived from fMRI using natural speech stimuli, where a semantic gradient spanned from perceptual and quantitative descriptions, as received from the external world, to human social interaction reflecting aspects of the internal self (41). The compression of this hierarchy in FEP may indicate a disturbance of these distinct levels of semantic processing and their relationship, possibly resulting in a decreased ability to differentiate the external world and the internal self.

Building higher-order meaning from perception is inherently subject to structure-building, over and above merely retrieving lexical-semantic concepts (42). In this respect, our behavioral linguistic analysis showed another surprising and previously undocumented pattern (Fig. 4A), which predicted the gradient dispersion of the semantic network: FEP exhibited *more* nodes in *higher* syntactic trees, and higher fraction scores of phrasal nodes out of word nodes, with the number of words as a covariate. This finding contrasts most previous studies (43), although Silva et al. (44) showed, in an overlapping sample, that while syntactic complexity dropped over time, it was higher at the FEP state. One possible explanation for the contrary findings in most prior studies, apart from illness stage, is that these might have been strongly biased by word count, whereby the schizophrenia group tended to produce shorter sentences probably with less syntactic complexity (21, 44–46). This motivated our inclusion of the number of words in the GLM model, though the FEP group generated broadly similar quantity of speech, as measured with words, utterances, and lexical categories, likely thanks to the control of speech duration within one minute. We suggest that the key to interpreting the result of greater syntactic depth in FEP is our finding of decreased approximate entropy of syntactic depth for each word in an utterance in FEP, in a context where the mean utterance *length* remained similar. Lower approximate entropy of syntactic depth suggests higher predictability of the syntactic depth of the next word given the previous words. The increased syntactic depth observed in FEP may thus have been achieved via a low-level process of simple word addition, rather than a genuine increase in hierarchical syntactic complexity. This would align with previous findings of less clausal embedding in psychosis (21, 45, 47, 48), a classical indicator of weakening hierarchical syntactic complexity.

Apart from higher syntactic tress and lower approximate entropy, the FEP group employed a greater number of verb phrases (VPs) to organize fewer, simpler, and shorter noun phrases (NPs). The intrinsic dispersion of the semantic brain network was predicted by the quantity of syntactic outputs, the hierarchical syntactic complexity, and predicate structures primarily associated with the VP, but not by argument structures as linked to the NP. This asymmetry between VPs and NPs matters, as they play different roles in the genesis of utterance-level meaning: VPs tend to encode *new* information, while NPs pick out objects/referents of which this new information is then predicated. New information generated about a referent can provide episodic details. Hence, a shift in the distribution of VPs and NPs may relate to shifts in episodic thinking processes mediated by semantic network regions.

### Limitations

Limitations firstly include that our dataset was relatively small, and we conducted several steps of analyses. Although we controlled for false discovery, type 2 errors cannot be fully ruled out. Furthermore, speech data were only collected in the context of a picture description task, which could potentially account for the observed correlation between semantic graph features and the dispersion of the first cortical gradient, rather than the dispersion of the semantic network gradient. Though it is a common practice to elicit spontaneous speech in such task, subsequent investigations should aim to validate our findings by employing speech elicited under various conditions on a larger dataset. In addition, it is worth noting that our research focused on the cortical structure and did not incorporate subcortical pathways, which could be vital in understanding psychosis (49).

### Conclusions

We demonstrated alterations of a unimodal-to-transmodal axis of cortical organization in first-episode psychosis and of a functional gradient in the semantic network. Both of these relate to the disintegration of meaning-making speech structure in patients. This confirms the role of spontaneous speech as one crucial and largely freely available behavioral readout of neurocognitive functioning as a whole. How we speak necessarily reflects how numerous cognitive processes are integrated at the level of the meaning we produce in response to a prompt or task: it reflects what we perceive and how we make conceptual sense of it, how we organize a discourse, dip into our semantic memory, and generate episodic information from concepts we remember. Psychosis involves large-scale shifts in the organization of neural cognitive substrates reflected in the process of generating meaning in language.

## Materials and Methods

### Dataset

Analyses were performed on twenty-nine untreated FEP patients recruited for this study, matched to twenty-nine healthy controls (HC) on age, gender, and education. The severity of symptoms in the patient group was assessed with the 8-item Positive and Negative Syndrome Scale (PANSS) version. The eight items are: delusions (PANSS8P1), conceptual disorganization (PANSS8P2), hallucinatory behavior (PANSS8P3), blunted affect (PANSS8N1), passive/apathetic social withdrawal (PANSS8N4), lack of spontaneity and flow of conversation (PANSS8N6), mannerisms & posturing (PANSS8G5), and unusual thought content (PANSS8G9). Subjects underwent a resting-state fMRI scan and were asked to describe three pictures from the Thematic Apperception Test (one minute for each picture) immediately prior to the scans. Demographic data with clinical scores are shown in Table 1. The whole workflow, including brain analyses and linguistic analyses, is shown in Figure 1. The patient group was recruited in the Prevention and Early Intervention Program for Psychosis in London, Ontario. Three psychiatrists made the clinical assessment for the diagnostic consensus, and the criteria were based on the Statistical Manual of Mental Disorders, 5^th^ Edition (DSM-5). At the time of testing, 50% of the patients were fully antipsychotic naïve, while the other 50% had an average antipsychotic exposure at the time of scan as 2.8 days (SD 3.7) while average the duration of untreated psychosis was 9.4 (SD 13.5) weeks.

### Resting-state *fMRI* acquisition and preprocessing

The fMRI data was acquired at the Centre for Functional and Metabolic Mapping (CFMM) at the University of Western Ontario on a Siemens 7 T Plus (Erlangen, Germany). A total of 360 whole-brain functional images were collected using a multi-band EPI acquisition sequence with 20 ms of echo time, 1000 ms of repetition time, a flip angle of 30◦, in 63 slices with a multi-band factor of 3, iPat of 3, and an isotropic resolution of 2 mm. The T1-weighted MP2RAGE anatomical volume was acquired at a 750 μm isotropic resolution (TE/TR = 2.83/6000 ms). Images were preprocessed using fMRIPrep 21.0.1 (50, 51) based on Nipype 1.6.1 (52, 53). Details on how fMRIPrep preprocessed the anatomical and functional data can be found in the supplementary information (SI-1).

### Cortical functional *hierarchy* with gradient dispersion and SFC

For the whole-brain gradient analysis, we extracted functional time-series from 1000 regions of interest (ROIs) using Schaefer 1000-parcels (30) (Fig. S1) organized into seven Yeo networks (54): visual (VN), somatomotor (SMN), dorsal attention (DAN), ventral attention (VAN), limbic (LN), frontoparietal control (FPN), and default mode (DMN). We computed a 1000×1000 FC matrix using z-transformed correlation coefficients of the time series. A standardized pipeline using the BrainSpace toolbox (31) was employed to map the FC profiles to low-dimensional fluids, and here we give a brief summary of the pipeline. We averaged the subject-level FC matrices for each group (HC and FEP) to compute a group-level template. The group templates were sparsified with all negative values, and 90% of the smallest elements per row were zeroed out to keep only positive, strong, and less noisy correlations. Then, for each two ROIs, we computed the normalized cosine similarity scores between their FC profiles for affinity matrices, and applied diffusion mapping to reduce the dimensionality to extract ten gradients. We also carried out this pipeline on every subject’s FC matrix. We aligned the subject-level gradient maps to the group-level gradient solution with generalized Procrustes rotation for more robustness. The first two gradients, G1 and G2, were analyzed due to their more established relationship with cognitive functions (55), with G1 reflecting the extent of separation between unimodal and transmodal cortex (i.e., from VN and SMN to DMN), and G2 differentiating SMN and VN (26).

Furthermore, we calculated the SFC based on the pipelines from Sepulcre et al. (13). The same FC matrices used for the gradient analysis were used to derive the statistical significance of correlation coefficients (SI-3.1), which were corrected with False Discovery Rate (FDR) for multiple comparisons (56), and binarized based on the corrected FDR *q*-values using the threshold of 0.001. The SFC degrees derived from such sparsified FC matrices indicate the count of all paths of a particular exact length, which connect a given seed region to all other areas (13). Following Sepulcre et al. (13), we used bilateral seeds of primary visual (V1), auditory (A1) and somatosensory (S1) areas (SI-3.2, Table S1 and Fig. S2-S3). While the gradient analyses shed light on connectivity segregation, SFC could serve as a window to the integration, which, from a small-wordless perspective, is not comprehended by investigating segregation only (57). SFC analysis starting in these unimodal areas has been shown to expand toward multimodal areas as the number of steps (i.e., the length of paths) grows, thus offering further insight into the first and second gradients. Similar to gradient analysis, we first computed the group-level SFC degrees from the group-level FC templates. SFC maps at every step were normalized using a min-max scaler into the range between 0 and 1. Secondly, for each subject, we computed the subject-level SFC degrees, normalized the map using the same scaler, and multiplied the subject-level SFC maps with the corresponding group-level SFC map at each step, which, as suggested by Hong et al. (58), increases the robustness of SFC maps. Spatial correlation across consecutive steps in group-level SFC maps, in both HC and FEP, converged at the sixth step (*r* > 0.999, Fig. S4-S5). We considered six steps to be sufficient for depicting connectivity profiles in our data.

### Functional hierarchy, gradient dispersion, and SFC in the semantic network

Within the semantic network, we first computed the ten gradients from FC matrices, analyzed the first gradient, and then defined the dispersion of the first gradient of the semantic network as the sum squared Euclidean distance of every node to its centroid. We manually matched Schaefer’s ROIs in the semantic network with these three subdivisions of the semantic network as grouped by Binder et al. (27), as in Table S4. Based on the gradient results (Table S3), where the hierarchy started from the heteromodal cortex and the multimodal and heteromodal association cortex to medial limbic regions, we chose two ROIs respectively for inferior frontal gyrus (IFG) and dorsolateral prefrontal cortex (dlPFC) as heteromodal seeds, and two ROIs for supramarginal gyrus (SMG) and angular gyrus (AG) as multimodal and heteromodal association seeds. The coordinates are available in Table S2 and Fig. S6. SFC maps in the semantic network converged at the fifth step (Fig. S7-S8).

### Graph-based analysis of semantic graphs in spontaneous speech

Semantic graphs at two levels, one of lexical categories and another of utterances, were constructed through analogous procedures. Tokenization and part-of-speech tagging were carried out with spaCy (en_core_web_sm, 3.4.2) (59). We computed the normalized cosine similarity scores between every pair of two embeddings, either lexical categories or utterances, for an affinity matrix binarized by proportional thresholding. Consistent with previous studies, we did not pick a single threshold in this case, as we did for gradient and SFC, but selected the lowest threshold from 0.05 to 0.8 with intervals of 0.05, which returns a sparsified matrix with the average degree (the degree of a node is the number of connections linked to the node) over all nodes larger than two multiplies the e-base logarithm of number of nodes (2 log N) (60). This procedure assures that the thresholded network exhibits small-world properties (61) with as few edges as possible. Notably, we did not require the small-worldness coefficients to be larger than 1.1 as done in some previous studies, as this is a value derived from specific brain graph data and did not apply to the semantic graph. In the next step, we moved from lexical categories to utterances as the basic units of narratives. The mean thresholds for lexical category graphs were 0.275 for the control group and 0.297 for FES, whereas the thresholds for utterance graphs were 0.448 for controls and 0.453 for FES. Group differences in the thresholds for both lexical category graphs (*p* = 0.170) and utterance graphs (*p* = 0.558) were not statistically significant, as indicated by GLMs with a Gaussian distribution and an identity link function. Age, education level, and sex were included as covariates in the GLMs. Semantic graphs were constructed for each narrative, instead of each utterance, because there were too few nodes in a considerable number of utterances to construct a meaningful graph (e.g., *I see a man and a woman*. There will be only three lexical categories *see*, *man*, and *woman* remained after removing stopwords, whereas at least four nodes are required for constructing a small-world graph (62).) All semantic measures were extracted from each of the three picture descriptions and averaged across them. Examples of the semantic graphs are provided in Fig. S10 (lexical categories) and Fig. S11 (utterances).

### Graph-based *analysis* of syntactic trees in spontaneous speech

Syntactic trees were derived from a constituency parser (benepar_en3) together with spaCy (59, 63). Punctuations were kept for parsing the trees but removed from the trees for subsequent analyses. The original trees from the parser were in Greibach normal form (GNF), which were converted to Chomsky normal form (CNF), allowing a parent node to have at most two children nodes, using the nltk package (3.7) (64). Trees were represented as directed acyclic graphs starting from the sentence node on top, through phrasal nodes, to the part-of-speech tags of every token forming the leaves of the tree at the bottom. Syntactic measures were extracted at the level of utterance. We first averaged the syntactic measures across utterance for each picture description, then averaged across picture descriptions for each subject. Examples of the syntactic trees are provided in Fig. S12.

### Statistical tests

The surface-based linear model (SLM) from the BrainStat toolbox (65) was employed to investigate the fixed effect of group on functional gradients and SFC degrees, with age, sex, and education level as covariates. *P*-values from SLM were corrected for multiple comparisons with false discovery rate (FDR). Corrected *p* values were reported as *q* values. Mann-Whitney U tests were carried out to compare the four gradient dispersion scores between HC and FEP, also corrected with FDR.

For language measures, we first correlated them with the quantity of speech production (lexical category and utterance counts) using partial Spearman’s ranked correlation with diagnosis as a covariate. Semantic graph measures were correlated with the number of lexical categories or utterances. Syntactic measures were correlated with the number of words. All results were domain-wise corrected for multiple comparisons using FDR. Specific classification of features into the domains can be found in SI-7. Then, we applied GLMs to explore the group differences of these language measures with age, sex, and education level as covariates. To regress out the effect of simply the quantity of speech production, the number of outputs was included as an additional covariate in those cases where correlations were significant. All semantic measures were compared with the number of lexical categories or utterances as the additional covariate (see Fig. S25). All syntactic measures, except for Phrase_fraction, were compared with the number of words as the additional covariate (see Fig. S26). We fit most of the data using Gaussian distribution with Identity link function, except for the number of words, where we used Tweedie distribution with log link function, and the numbers of entities and utterances, where Gamma distribution with log link function was used. Deviance goodness-of-fit tests indicated all GLMs fit the data well (*p* > 0.05).

To explore the relationship between functional hierarchy and different language measures, we employed GLM models with Tweedie distribution and log link function to predict the four gradient dispersion scores as descriptors of the functional hierarchy.

All statistical tests were carried out with statsmodels (0.13.5) and pingouin (0.5.3). Statistical significance was acknowledged when the *p-*value (or *q-*value when corrected) was less than 0.05.

## Data and code availability

All codes for analysis were developed by the authors in Python (3.9.12) using free and open-source packages. All scripts will be available at https://github.com/RuiHe1999/FEP_gradient_sem_syn.

## Acknowledgments

We appreciate all the participants and their families for the time and effort to contribute to this study. We also acknowledge Michael Mackinley, Jenny Chan, and Sabrina Ford of Western University for their support in preparing the dataset. This research was supported by European Research Council (ERC-2023-SyG, 101118756 to WH), China Scholarship Council (grant 202108390062 to RH), the Department of Science and Technology of Guangdong Province (grant 112175605105 to WH and RH), and the National Agency for Research and Development (ANID), Scholarship Program, Becas Chile 2019, Postdoctoral Fellow 74200048 (MA) (to MFAS). The data acquisition for this study was funded by CIHR Foundation Grant (FDN 154296) to LP and was supported by the Canada First Excellence Research Fund to BrainSCAN, Western University (Imaging Core); Innovation fund for Academic Medical Organization of Southwest Ontario; Bucke Family Fund, The Chrysalis Foundation, The Children Hospital Foundation and The Arcangelo Rea Family Foundation (London, Ontario). Compute Canada Resources (Application No. 1530) were used in the storage and analysis of imaging data. LP acknowledges research support from the Canada First Research Excellence Fund, awarded to the Healthy Brains, Healthy Lives initiative at McGill University (New Investigator Supplement); Monique H. Bourgeois Chair in Developmental Disorders and Graham Boeckh Foundation (Douglas Research Centre, McGill University) and a salary award from the Fonds de recherche du Quebec-Sante (FRQS).

