## Supplementary material for "Changes in the structure of spontaneous speech predict the disruption of hierarchical brain organization in first-episode psychosis": SI Appendix

### 1. Information on reprocessing using fMRIPrep

**Anatomical data preprocessing.** A total of 1 T1-weighted (T1w) images were found within the input BIDS dataset. The T1-weighted (T1w) image was corrected for intensity non-uniformity (INU) with N4BiasFieldCorrection(1), distributed with ANTs 2.3.3(2), and used as T1w-reference throughout the workflow. The T1w-reference was then skull-stripped with a Nipype implementation of the antsBrainExtraction.sh workflow (from ANTs), using OASIS30ANTs as target template. Brain tissue segmentation of cerebrospinal fluid (CSF), white-matter (WM) and gray-matter (GM) was performed on the brain-extracted T1w using fast (FSL 6.0.5.1:57b01774, RRID:SCR_002823)(3). Brain surfaces were reconstructed using recon-all (FreeSurfer 6.0.1, RRID:SCR_001847)(4), and the brain mask estimated previously was refined with a custom variation of the method to reconcile ANTs-derived and FreeSurfer-derived segmentations of the cortical gray-matter of Mindboggle (RRID:SCR_002438)(5). Volume-based spatial normalization to one standard space (MNI152NLin2009cAsym) was performed through nonlinear registration with antsRegistration (ANTs 2.3.3), using brain-extracted versions of both T1w reference and the T1w template. The following template was selected for spatial normalization: ICBM 152 Nonlinear Asymmetrical template version 2009c [RRID:SCR_008796; TemplateFlow ID: MNI152NLin2009cAsym](6).

**Functional data preprocessing.** For each of the 1 BOLD runs found per subject (across all tasks and sessions), the following preprocessing was performed. First, a reference volume and its skull-stripped version were generated using a custom methodology of fMRIPrep. Head-motion parameters with respect to the BOLD reference (transformation matrices, and six corresponding rotation and translation parameters) are estimated before any spatiotemporal filtering using mcflirt (FSL 6.0.5.1:57b01774)(7). The BOLD time-series (including slice-timing correction when applied) were resampled onto their original, native space by applying the transforms to correct for head-motion. These resampled BOLD time-series will be referred to as preprocessed BOLD in original space, or just preprocessed BOLD. The BOLD reference was then co-registered to the T1w reference using bbregister (FreeSurfer) which implements boundary-based registration(8). Co-registration was configured with six degrees of freedom. Several confounding time-series were calculated based on the preprocessed BOLD: framewise displacement (FD), DVARS and three region-wise global signals. FD was computed using two formulations following Power (absolute sum of relative motions)(9) and Jenkinson (relative root mean square displacement between affines)(7). FD and DVARS are calculated for each functional run, both using their implementations in Nipype (following the definitions by Power et al.(9)). The three global signals are extracted within the CSF, the WM, and the whole-brain masks. Additionally, a set of physiological regressors were extracted to allow for component-based noise correction (CompCor)(10). Principal components are estimated after high-pass filtering the preprocessed BOLD time-series (using a discrete cosine filter with 128s cut-off) for the two CompCor variants: temporal (tCompCor) and anatomical (aCompCor). tCompCor components are then calculated from the top 2% variable voxels within the brain mask. For aCompCor, three probabilistic masks (CSF, WM and combined CSF+WM) are generated in anatomical space. The implementation differs from that of Behzadi et al.(10) in that instead of eroding the masks by 2 pixels on BOLD space, the aCompCor masks are subtracted a mask of pixels that likely contain a volume fraction of GM. This mask is obtained by dilating a GM mask extracted from the FreeSurfer's aseg segmentation, and it ensures components are not extracted from voxels containing a minimal fraction of GM. Finally, these masks are resampled into BOLD space and binarized by thresholding at 0.99 (as in the original implementation). Components are also calculated separately within the WM and CSF masks. For each CompCor decomposition, the k components with the largest singular values are retained, such that the retained components' time series are sufficient to explain 50 percent of variance across the nuisance mask (CSF, WM, combined, or temporal). The remaining components are dropped from consideration. The head-motion estimates calculated in the correction step were also placed within the corresponding confounds file. The confound time series derived from head motion estimates and global signals were expanded with the inclusion of temporal derivatives and quadratic terms for each.(11) Frames that exceeded a threshold of 0.5 mm FD or 1.5 standardised DVARS were annotated as motion outliers. The BOLD time-series were resampled into standard space, generating a preprocessed BOLD run in MNI152NLin2009cAsym space. First, a reference volume and its skull-stripped version were generated using a custom methodology of fMRIPrep. All resamplings can be performed with a single interpolation step by composing all the pertinent transformations (i.e. head-motion transform matrices, susceptibility distortion correction when available, and co-registrations to anatomical and output spaces). Gridded (volumetric) resamplings were performed using antsApplyTransforms (ANTs), configured with Lanczos interpolation to minimize the smoothing effects of other kernels.(12) Non-gridded (surface) resamplings were performed using mri_vol2surf (FreeSurfer).

### 2. Shaefer parcellation

Below we visualized Schaefer's 1000-parcels (7 networks, resolution of 2 mm) (13), different colors indicating the seven Yeo's network: visual network (VN), somatomotor network (SMN), dorsal attention network (DAN), ventral attention network (VAN), limbic network (LN), frontoparietal control network (FPN), and default mode network (DMN) (14). Binder's semantic network greatly overlapped with the left DMN and LN. Visualization is shown in Fig. S1.

### 3. Additional methodological details of step-wise functional connectivity in cortex

**Compute FC matrix with p and q values for SFC analysis.** We utilized a function which could derive significant levels for Pearson correlation based on the correlation coefficient and degree of freedom, publicly available at: https://stackoverflow.com/a/24547964. The function is:

```
import numpy as np
from scipy.special import betainc

def corrcoef(matrix):
    r = np.corrcoef(matrix)
    rf = r[np.triu_indices(r.shape[0], 1)]
    df = matrix.shape[1] - 2
    ts = rf * rf * (df / (1 - rf * rf))
    pf = betainc(0.5 * df, 0.5, df / (df + ts))
    p = np.zeros(shape=r.shape)
    p[np.triu_indices(p.shape[0], 1)] = pf
    p[np.tril_indices(p.shape[0], -1)] = p.T[np.tril_indices(p.shape[0], -1)]
    p[np.diag_indices(p.shape[0])] = np.ones(p.shape[0])
    return r, p
```

We tested this function against the iterative usage of Pearson correlation from scipy.stats:

```
from scipy.stats import pearsonr

def corrcoef_loop(matrix):
    rows, cols = matrix.shape[0], matrix.shape[1]
    r = np.ones(shape=(rows, rows))
    p = np.ones(shape=(rows, rows))
    for i in range(rows):
        for j in range(i+1, rows):
            r_, p_ = pearsonr(matrix[i], matrix[j])
            r[i, j] = r[j, i] = r_
            p[i, j] = p[j, i] = p_
    return r, p
```

The coefficient and *p*-value matrices returned by these two functions are identical, tested on a randomly generated array (np.random.randn(1000, 359), same shape as the time series). The first function, which we used, brought two benefits: (1) being more efficient by almost 1300 times (3.21s for the *corrcoef* function while 4115.48s for *corrcoef_loop* function); (2) allowing us to compute significance matrix without the need of original time series.

**Visualization of SFC ROIs.** The coordinates of ROIs for step-wise functional connectivity (SFC) and corresponding name in Schafer's parcellation were listed in Table S1 and visualized in Fig. S2. LH for left hemisphere while RH for right hemisphere. The ROIs were seeds for the primary visual (V1), auditory (A1) and somatosensory (S1) areas. MNI coordinates was transformed to the name of ROIs in Schafer's parcellation by FSLeyes (FSLVM7_64), as in Fig. S3.

**Spatial correlation to decide the number of steps to report.** In line with Sepulcre et al.(15), we computed the Pearson correlation between every two consecutive steps of the three seed-based SFC maps for the first ten steps, and recognized a convergence when correlation coefficient is larger than 0.999. Such spatial correlation suggested the SFC maps became stable after the sixth step, as shown in Fig. S4 and Fig. S5.

### 4. Additional methodological details of SFC in semantic network

The coordinates of ROIs for step-wise functional connectivity (SFC) and corresponding names in Schafer's parcellation, AAL3, and Broadman were listed in Table S2 and visualized in Fig. S6. Spatial correlations showed the convergence at the fifth step, as in Fig. S7 and Fig. S8.

### 5. The list of stop-words

['i', 'me', 'my', 'myself', 'we', 'our', 'ours', 'ourselves', 'you', "you're", "you've", "you'll", "you'd", 'your', 'yours', 'yourself', 'yourselves', 'he', 'him', 'his', 'himself', 'she', "she's", 'her', 'hers', 'herself', 'it', "it's", 'its', 'itself', 'they', 'them', 'their', 'theirs', 'themselves', 'what', 'which', 'who', 'whom', 'this', 'that', "that'll", 'these', 'those', 'am', 'is', 'are', 'was', 'were', 'be', 'been', 'being', 'have', 'has', 'had', 'having', 'do', 'does', 'did', 'doing', 'a', 'an', 'the', 'and', 'but', 'if', 'or', 'because', 'as', 'until', 'while', 'of', 'at', 'by', 'for', 'with', 'about', 'against', 'between', 'into', 'through', 'during', 'before', 'after', 'above', 'below', 'to', 'from', 'up', 'down', 'in', 'out', 'on', 'off', 'over', 'under', 'again', 'further', 'then', 'once', 'here', 'there', 'when', 'where', 'why', 'how', 'all', 'any', 'both', 'each', 'few', 'more', 'most', 'other', 'some', 'such', 'no', 'nor', 'not', 'only', 'own', 'same', 'so', 'than', 'too', 'very', 's', 't', "'s", "'t", "'d", "'m", "'ve", "'ll", "'re", "'o", "'y", "'ain", 'can', 'will', 'just', 'don', "don't", 'should', "should've", 'now', 'd', 'll', 'm', 'o', 're', 've', 'y', 'ain', 'aren', "aren't", 'could', 'couldn', "couldn't", 'didn', "didn't", 'doesn', "doesn't", 'hadn', "hadn't", 'hasn', "hasn't", 'haven', "haven't", 'isn', "isn't", 'ma', 'might', 'mightn', "mightn't", 'must', 'mustn', "mustn't", 'need', 'needn', "needn't", 'shall', 'shan', "shan't", 'should', 'shouldn', "shouldn't", 'wasn', "wasn't", 'weren', "weren't", 'will', 'won', "won't", 'would', 'wouldn', "wouldn't", "n't", "um", "umm", "uh", "uhh", "yeah", "oh", "ah", "okay", "hm", "mhmm", "hmm", "hmmm", "well", "alright"]

### 6. Examples of language graphs

This is an example description of the field picture described by one of the authors (one-minute spontaneous speech) and divided into utterances:

- *It seems to be a picture about life on farm .*
- *A man is holding his horse in the background .*
- *It seems that they are gonna plough the fields .*
- *The woman is leaning onto the tree in the left side .*
- *She's holding her abdomen .*
- *Seems like she's pregnant .*
- *On the right side, there's a girl holding a book, looking to the road .*
- *Maybe she's waiting for someone .*
- *The farm locates in a mountain area .*
- *Seems like in a village, as there are other houses in the background .*

Threshold for the semantic similarity using FastText (FT) embeddings was 0.25 and the generated graph was illustrated as in Fig. S9. Threshold for the semantic similarity using sentence-transformers (ST) embeddings was 0.55 and the generated graph was illustrated as in Fig. S10.

In addition, we generated a syntactic graph for the sentence: *On the right side, there's a girl holding a book, looking to the road .* The syntactic tree was shown in Fig. S11.

### 7. Domains for FDR correction

Syntactic:
Quantity of outputs: ['Leaf_num'],
Hierarchical complexity: ['Depth', 'Td'],
Quantity of production: ['Nodes', 'Phrase_fraction'],
NP-related measures: ['NP_count', 'NP_nest', 'NP_length'],
VP-related measures: ['VP_count', 'VP_nest', 'VP_length'],

Semantic:
Quantity of outputs: ['Ent_Num', 'Utt_Num', ],
Semantic similarity: ['FT_Sim', 'ST_Sim', ],
FT-graph measures: ['FT_threshold', 'FT_CC', 'FT_GE', 'FT_Clustering', 'FT_Sigma'],
ST-graph measures: ['ST_threshold', 'ST_CC', 'ST_GE', 'ST_Clustering', 'ST_Sigma']

Gradient dispersion: ['VN_DMN', 'SMN_DMN', 'VN_SMN', 'SemN_G1']

### 8. Group templates of functional gradient

**Whole cortex.** Group-level cortical gradient templates from the principal gradient to the tenth gradient were visualized as shown in Fig. S12 for HC and Fig. S13 for FEP. Scree plot showing the eigenvalue of each gradient were visualized as shown in Fig. S14 for HC and Fig. S15 for FEP.

**Semantic network.** Group-level semantic network gradient templates were visualized as shown in Fig. S16 for HC and Fig. S17 for FEP. Scree plot showing the eigenvalue of each gradient were visualized as shown in Fig. S18 for HC and Fig. S19 for FEP.

The ROIs and their RAS coordinates were downloadable from https://github.com/ThomasYeoLab/CBIG/blob/master/stable_projects/brain_parcellation/Schaefer2018_LocalGlobal/Parcellations/MNI/Centroid_coordinates/Schaefer2018_1000Parcels_7Networks_order_FSLMNI152_2mm.Centroid_RAS.csv. Below we gave the five ROIs with the smallest five gradient values and the largest five gradient values, their RAS coordinates (of the centroids, the RAS coordinates were taken as the MNI coordinates as the parcellation is in a standard MNI152 space), and the Brodmann Areas (BA) where the coordinates located (BA: https://bioimagesuiteweb.github.io/webapp/mni2tal.html) as in Table S3. The whole table of ROIs, RAS coordinates, the corresponding cortices, and BA areas were available in Table S4.

### 9. Group templates of SFC

**Whole cortex.** Group-level SFC maps from the first to the sixth steps were visualized as shown in Fig. S20 for the visual seed, Fig. S21 for the somatosensory seed, and Fig. S22 for the auditory seed.

**Semantic network.** Group-level SFC maps within the semantic network from the first to the fifth steps were visualized as shown in Fig. S23 and Fig. S24.

### 10. Correlation between linguistic measures and the quantity of speech

Correlations were visualized as shown in Fig. S25 for semantic analyses (the number of lexical categories or utterances) and Fig. S26 for syntactic analyses (the number of words).

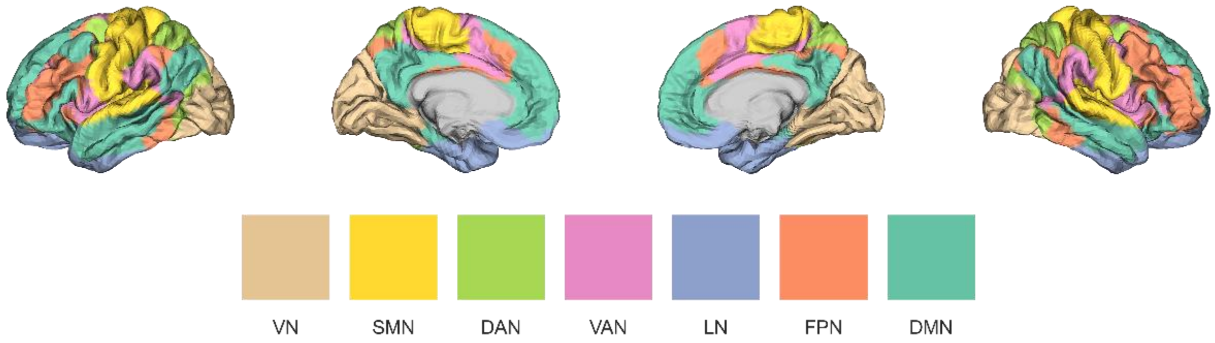


**Fig. S1.** Visualization of Schafer's 1000-parcels.

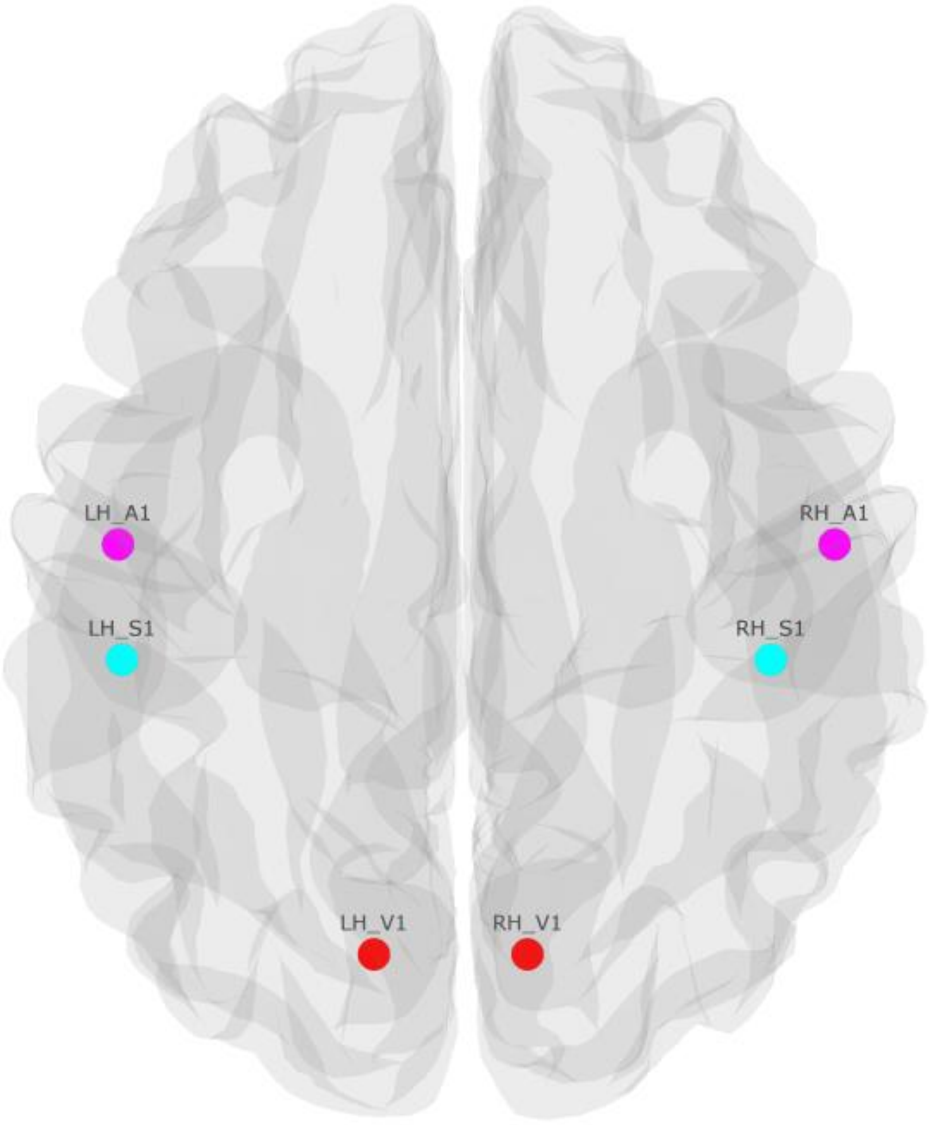


**Fig. S2.** SFC ROIs (top view), red: V1, magenta: A1, cyan: S1.

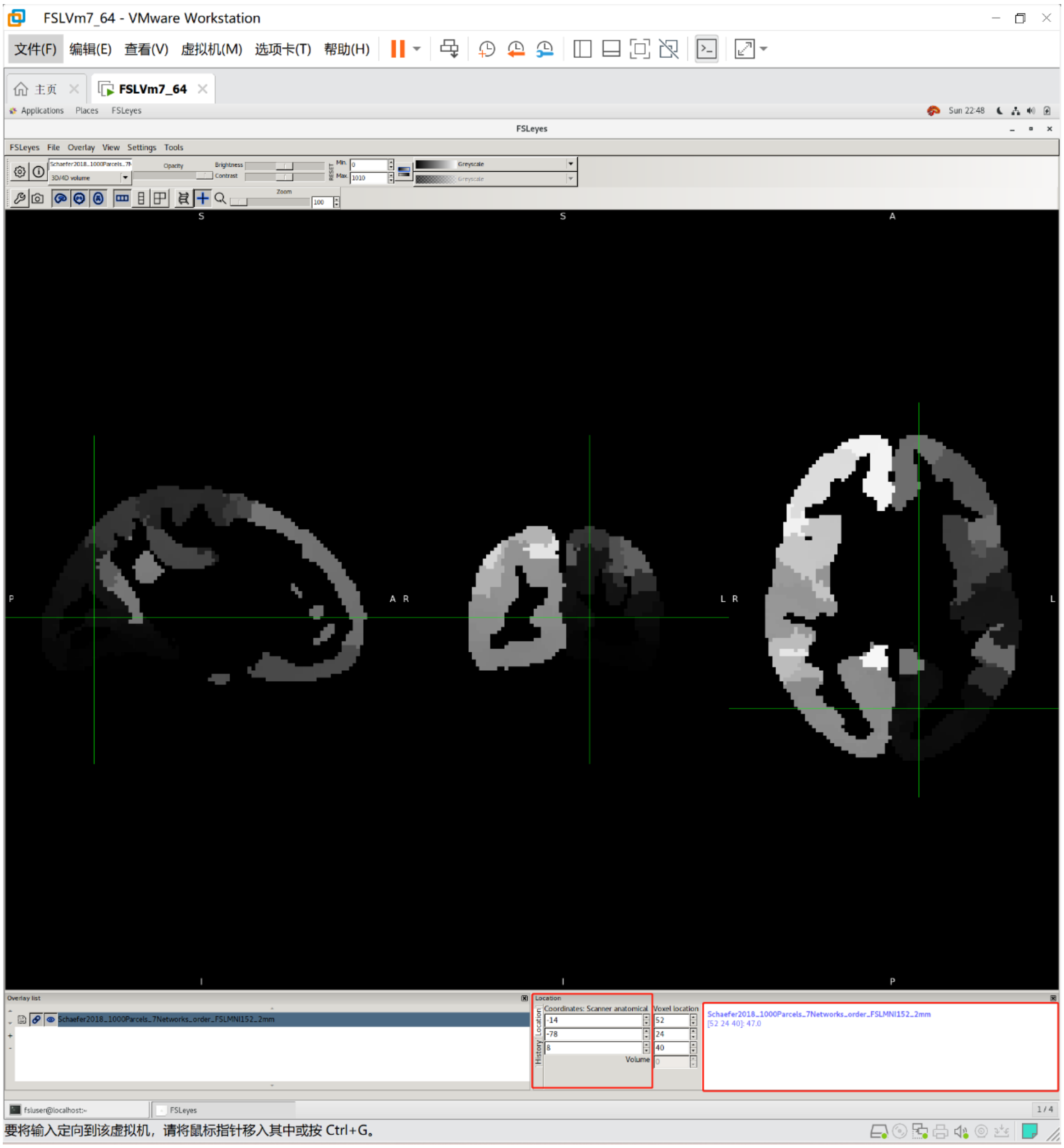


**Fig. S3.** Example of translating the MNI coordinate (-14, -78, 8) to name of ROIs in the parcellation. As shown in the bottom-rightmost panels at bottom, this coordinate locates at the 47th ROI, i.e. 7Networks_LH_Vis_47.

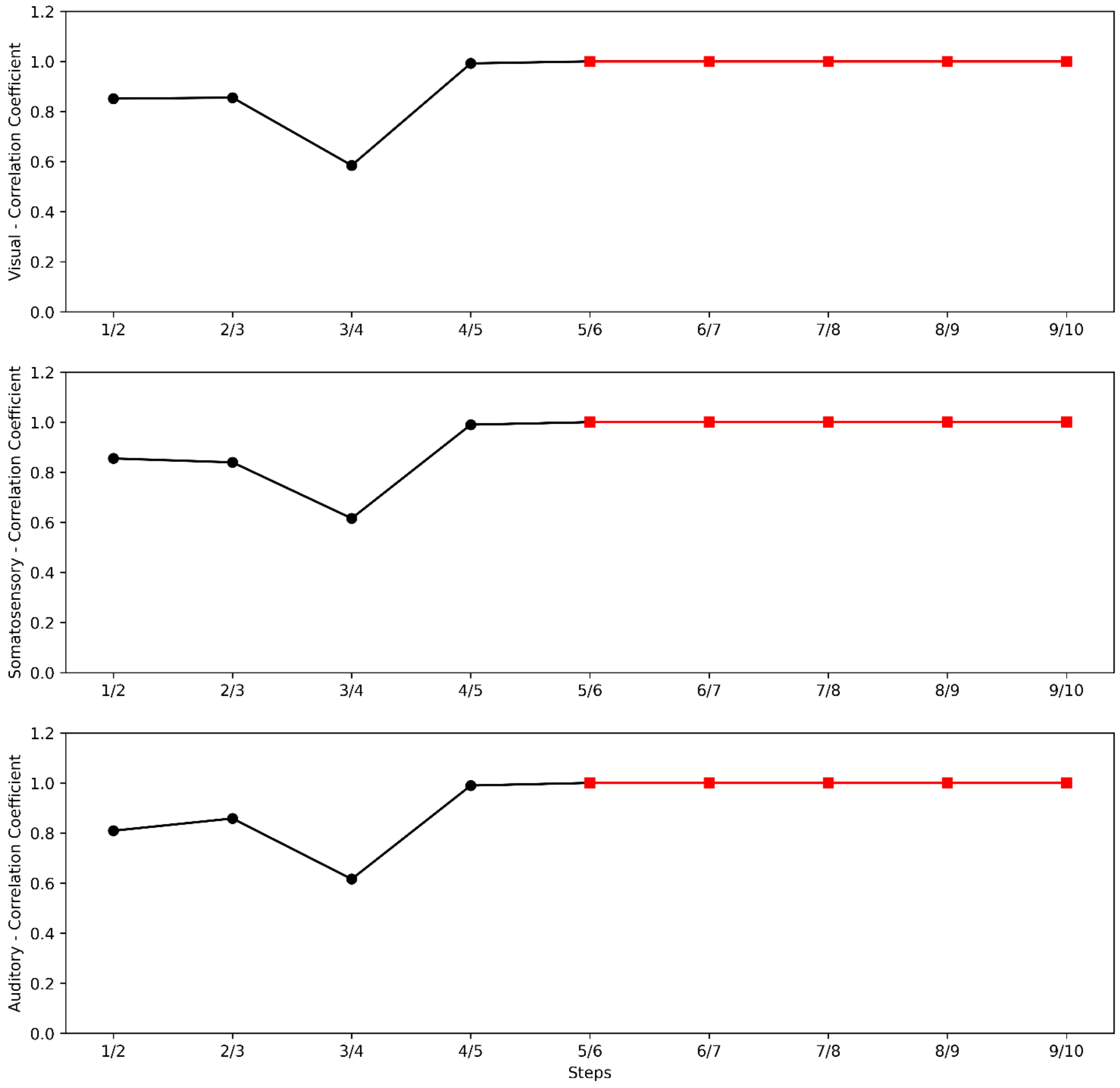


**Fig. S4.** Spatial correlation across steps in HC, from the first/second to the ninth/tenth. The results showed stable correlation coefficients after a five to six link-step distance (red squares and lines), defined as the correlation coefficients larger than 0.999.

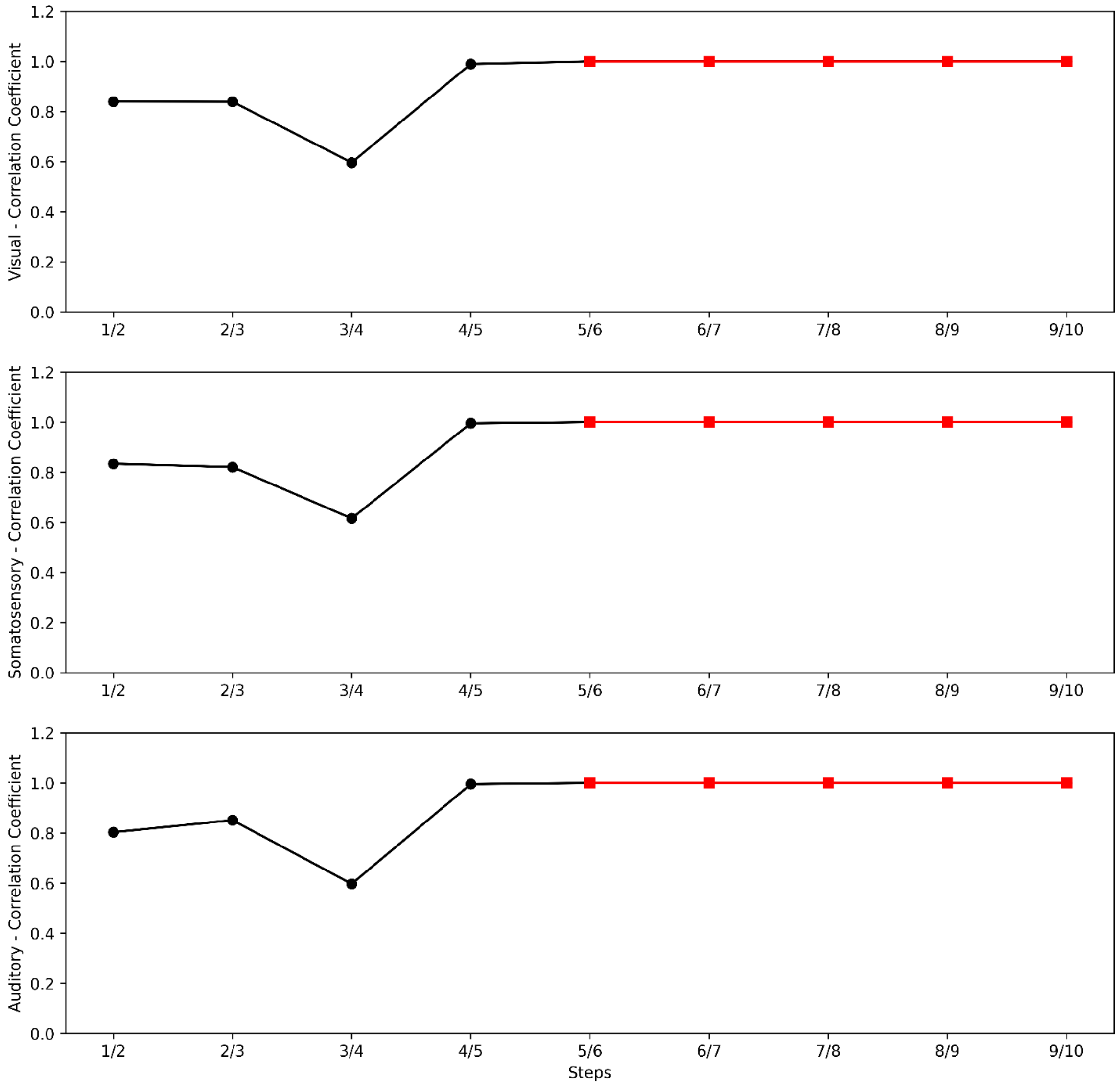


**Fig. S5.** Spatial correlation across steps in FEP, from the first/second to the ninth/tenth. The results showed stable correlation coefficients after a five to six link-step distance (red triangles and lines), defined as the correlation coefficients larger than 0.999.

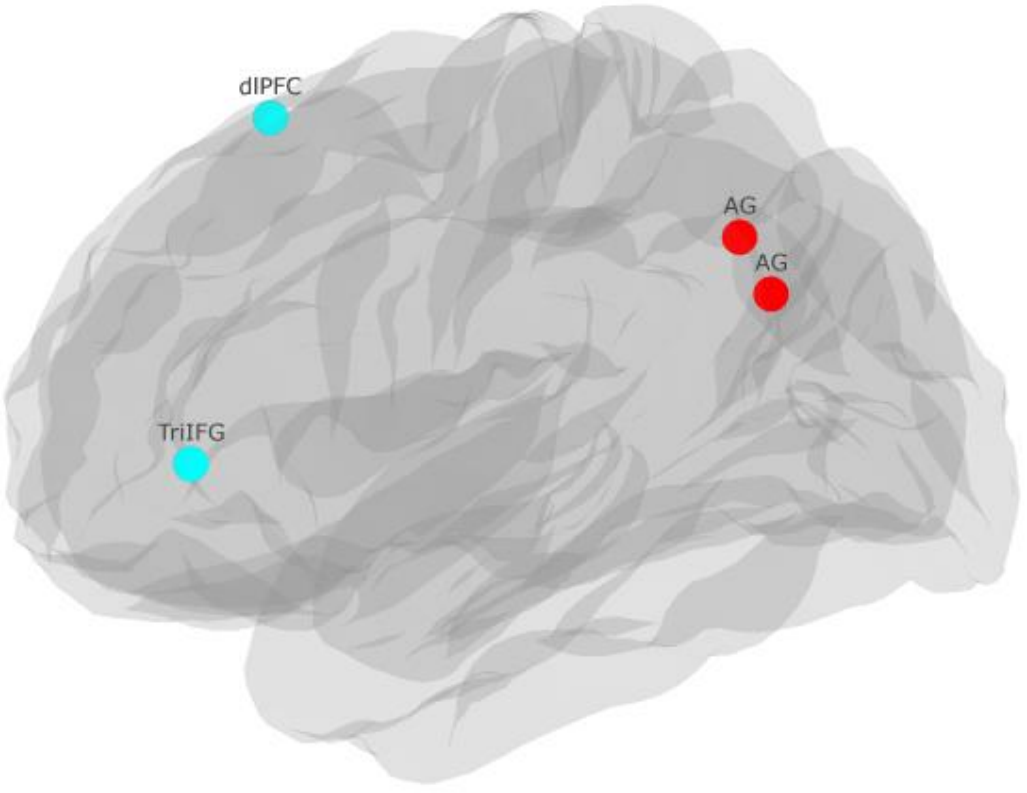


**Fig. S6.** SFC ROIs (left view). Blue: frontal heteromodal association seeds in the left hemisphere; red: parietal heteromodal association seeds in the left hemisphere.

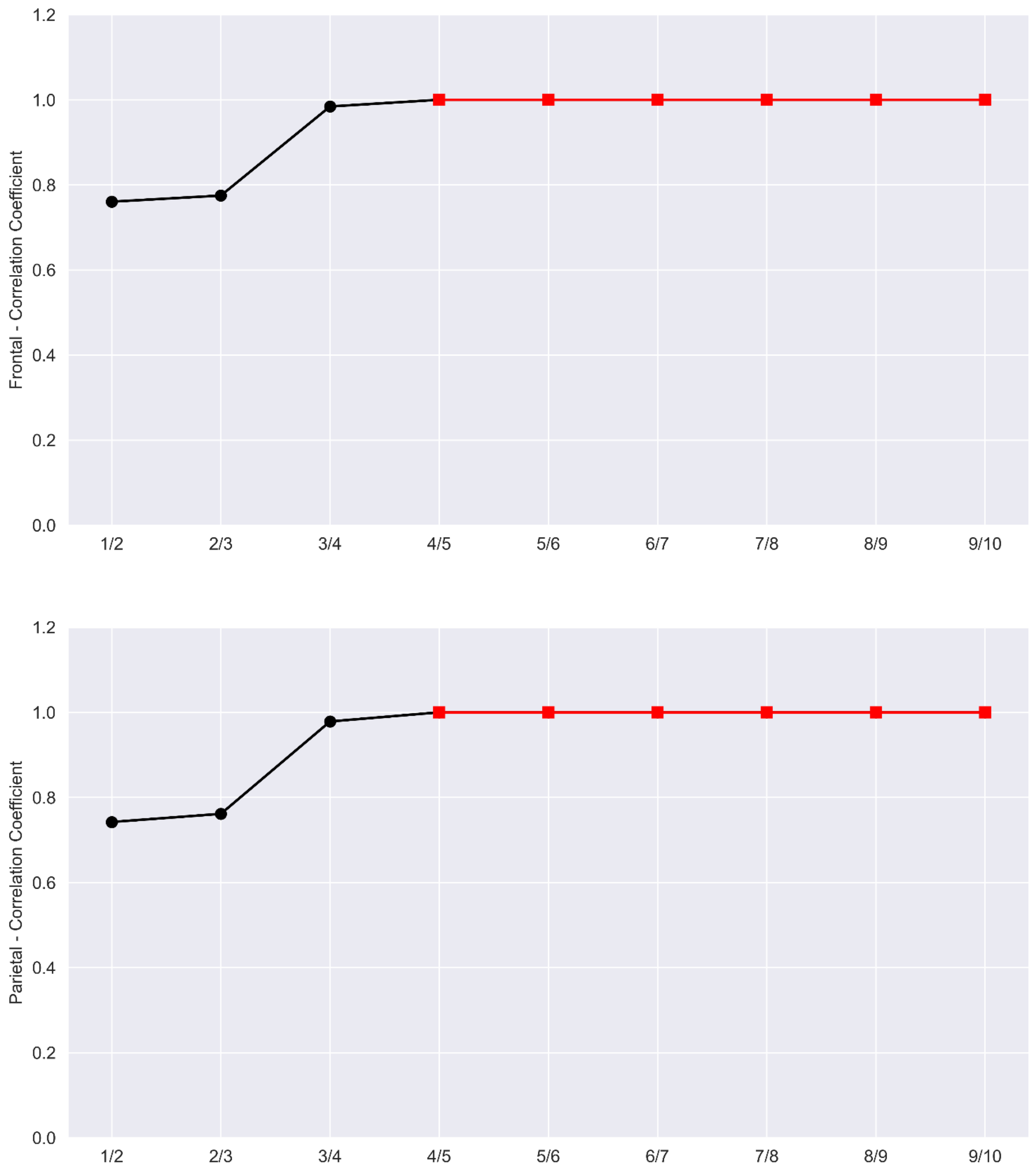


**Fig. S7.** Spatial correlation across steps in HC, from the first/second to the ninth/tenth. The results showed stable correlation coefficients after a four to five link-step distance (red triangles and lines), defined as the correlation coefficients larger than 0.999.

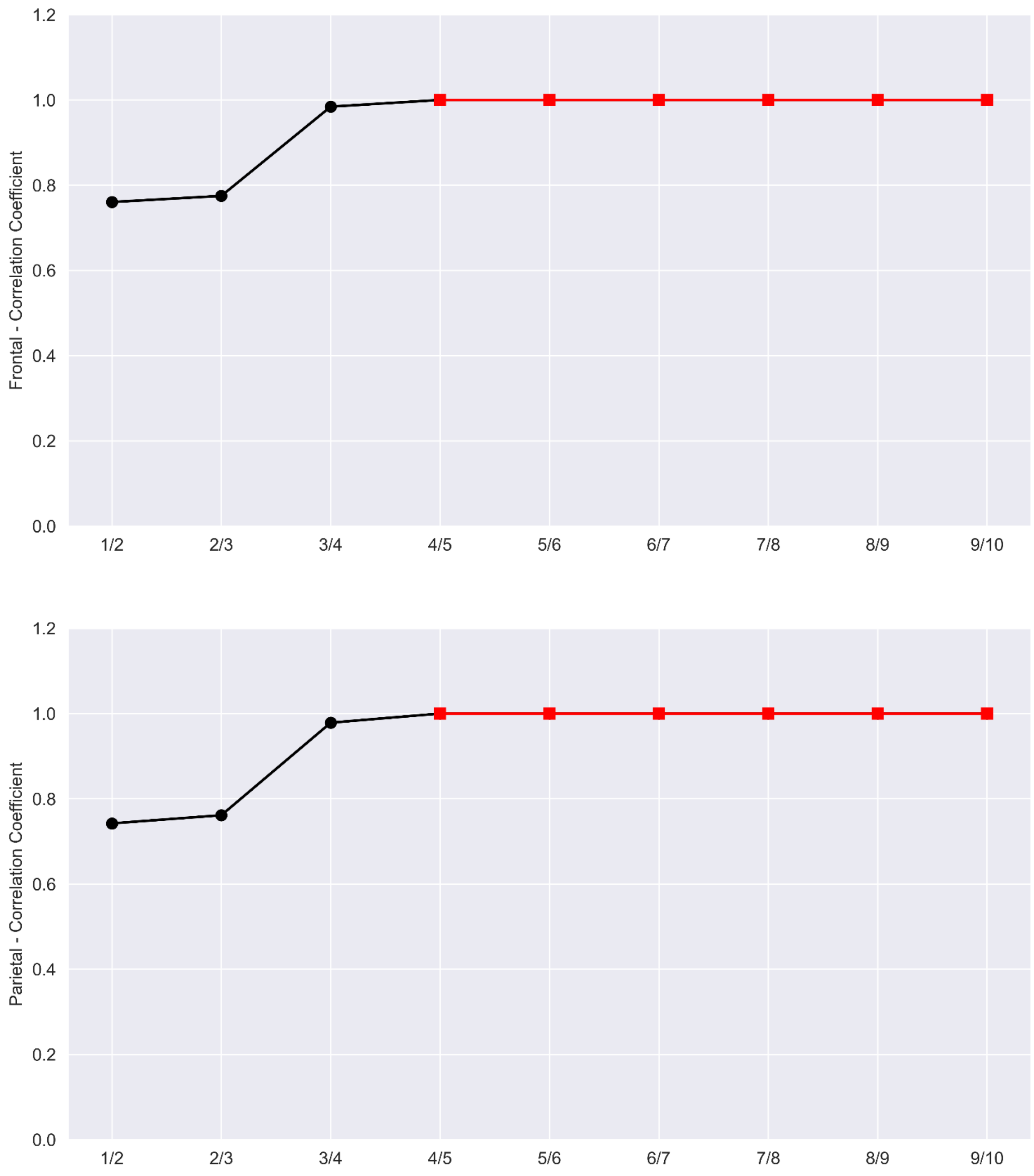


**Fig. S8.** Spatial correlation across steps in FEP, from the first/second to the ninth/tenth. The results showed stable correlation coefficients after a four to five link-step distance (red triangles and lines), defined as the correlation coefficients larger than 0.999.

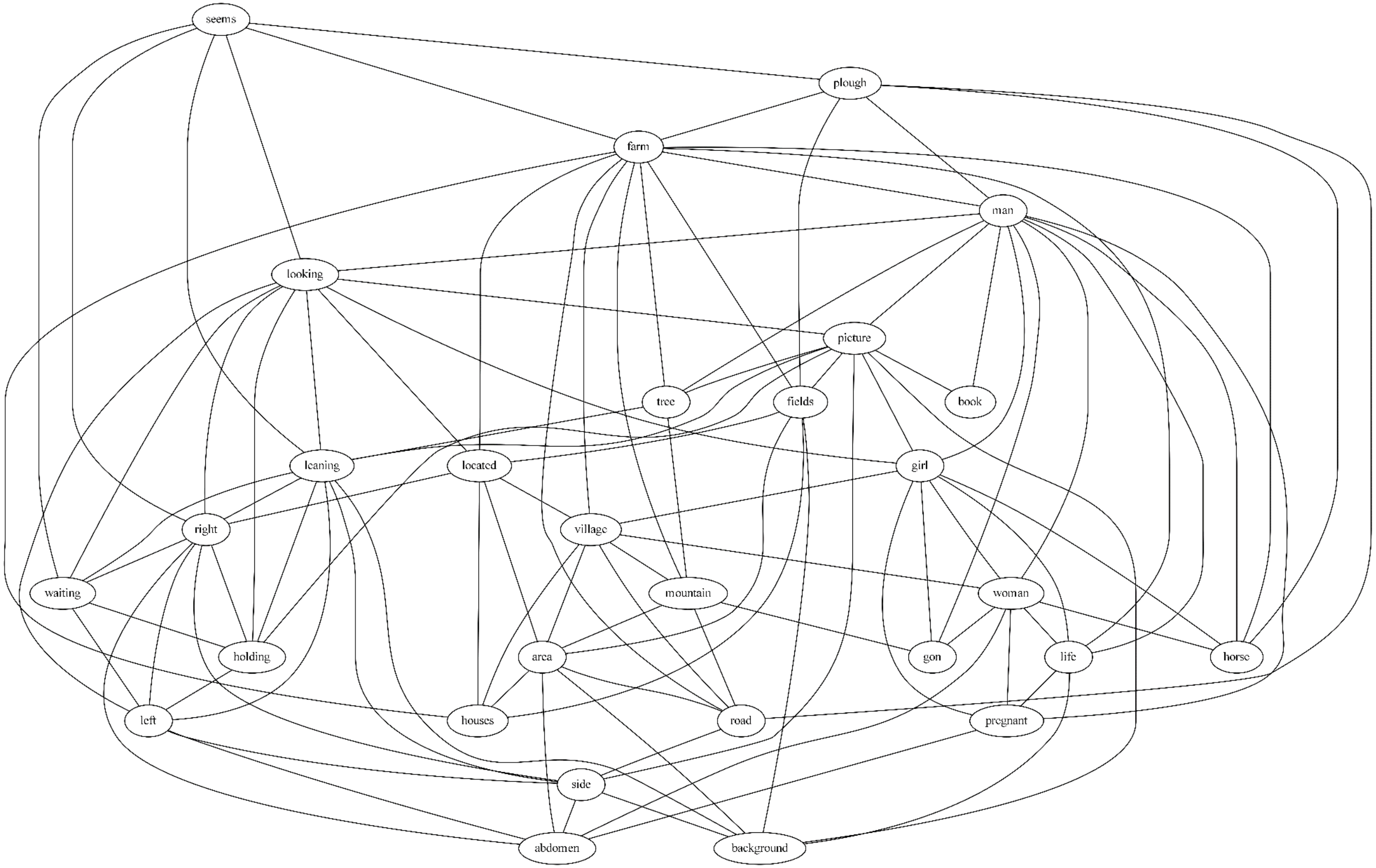


**Fig. S9.** Lexical category graph based on FT embeddings. Labels are lexical entities extracted from the narrative.

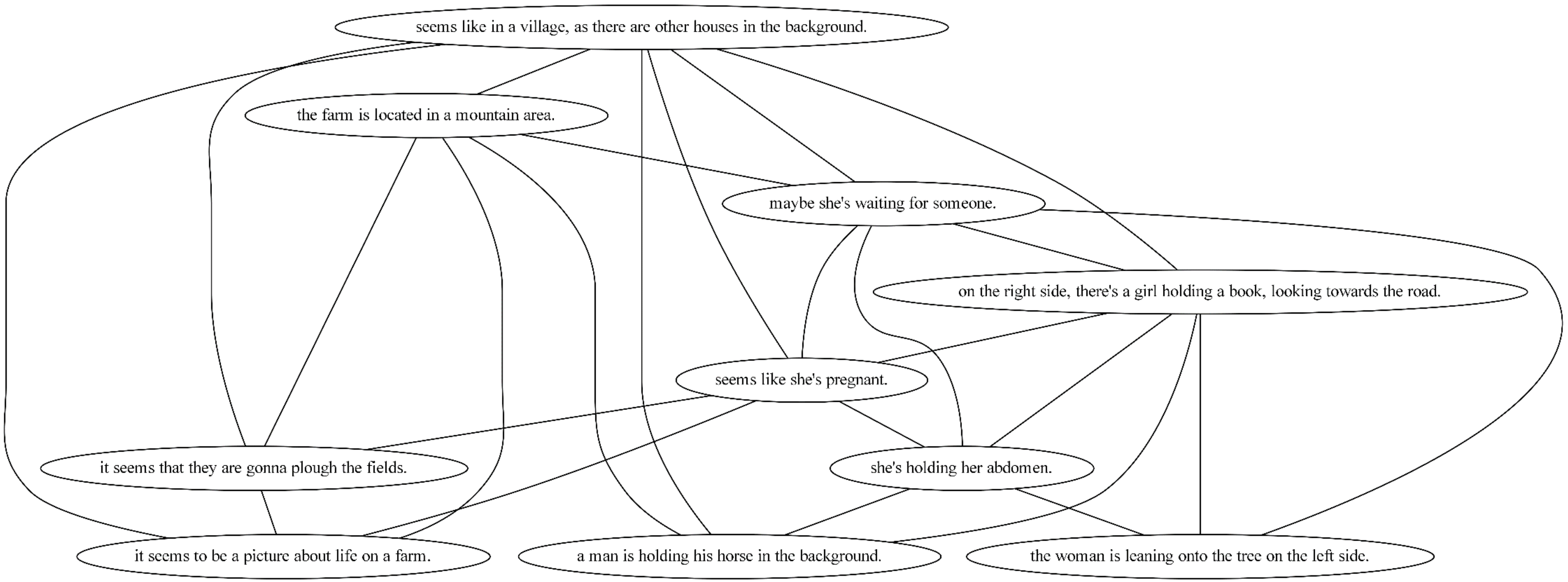


**Fig. S10.** Sentence graph based on ST embeddings. Labels are utterances.

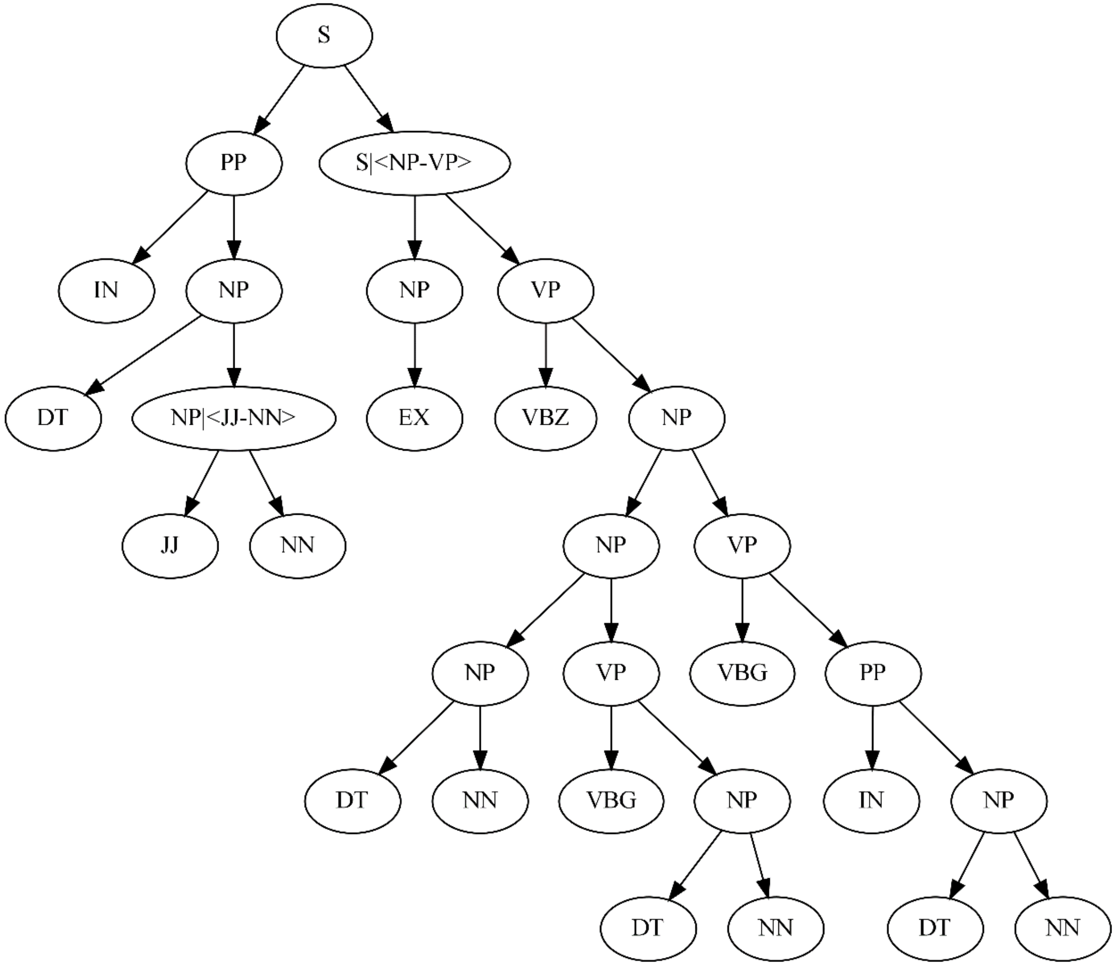


**Fig. S11.** Syntactic tree (as a directed acyclic graph) in Chomsky Norm Form. The leaf nodes are, from left to right: ['on', 'the', 'right', 'side', 'there', "'s", 'a', 'girl', 'holding', 'a', 'book', 'looking', 'to', 'the', 'road']. The abbreviations are: DT: determiner; EX: Existential there; IN: Preposition or subordinating conjunction; JJ: Adjective; NN: Noun, singular or mass; NP: noun phrase; PP: prepositional phrase; S: sentence; VBG: verb, gerund or present participle; VBZ: verb, 3rd person singular present; VP: verb phrase.

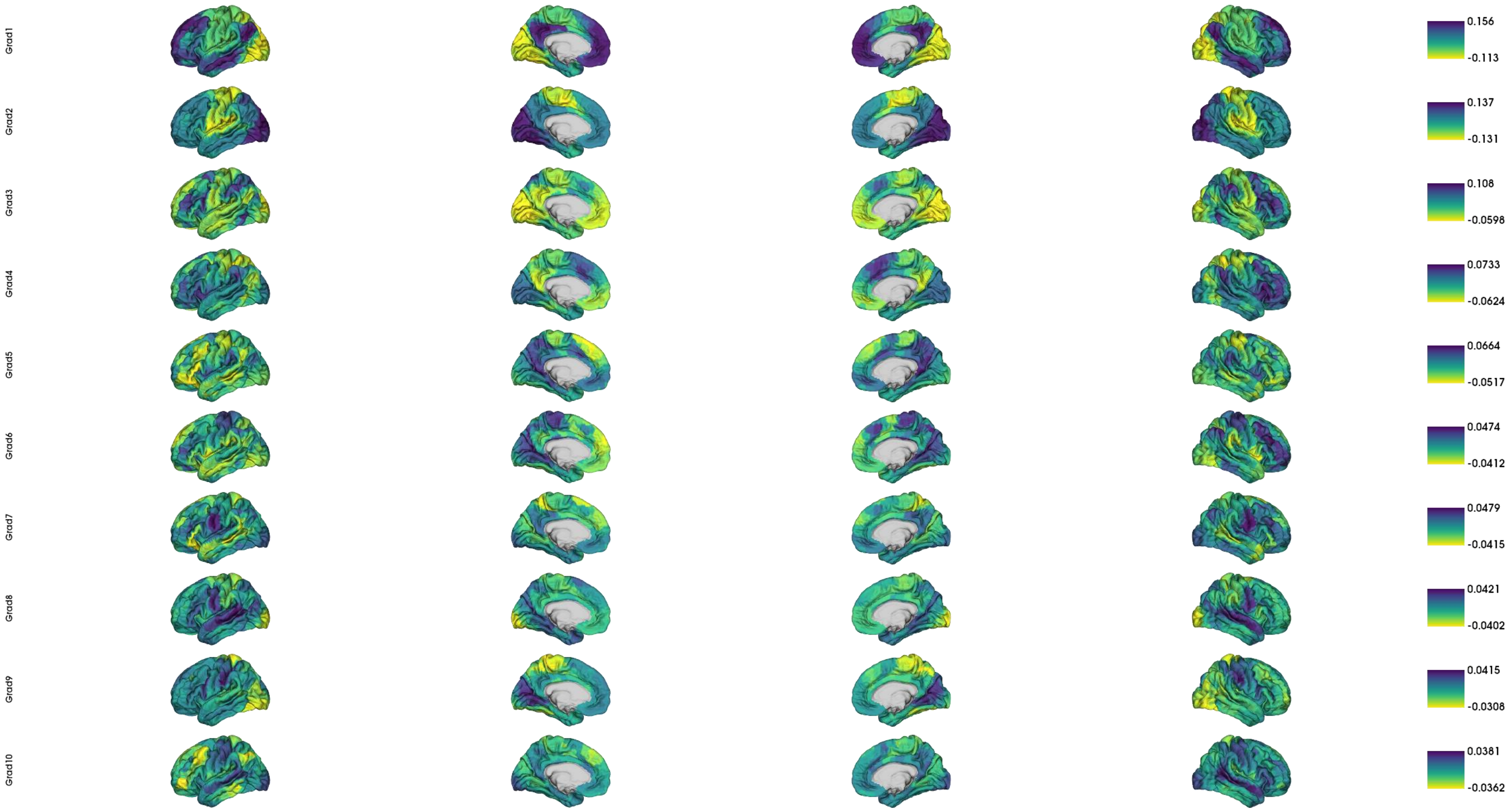


**Fig. S12.** Group templates of cortical gradients from the principal gradient to the tenth gradient in the HC group.

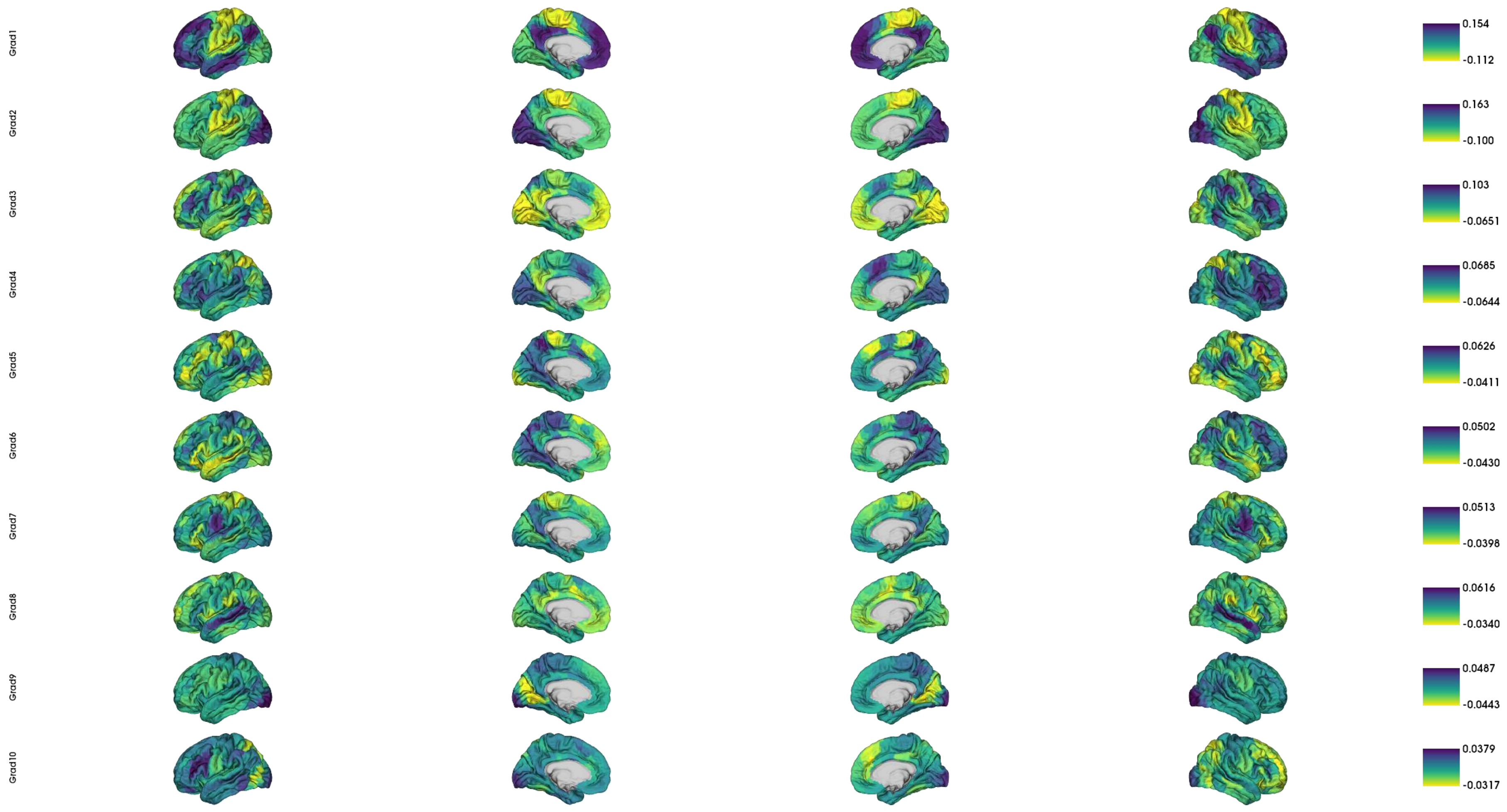


**Fig. S13.** Group templates of cortical gradients from the principal gradient to the tenth gradient in the FES group.

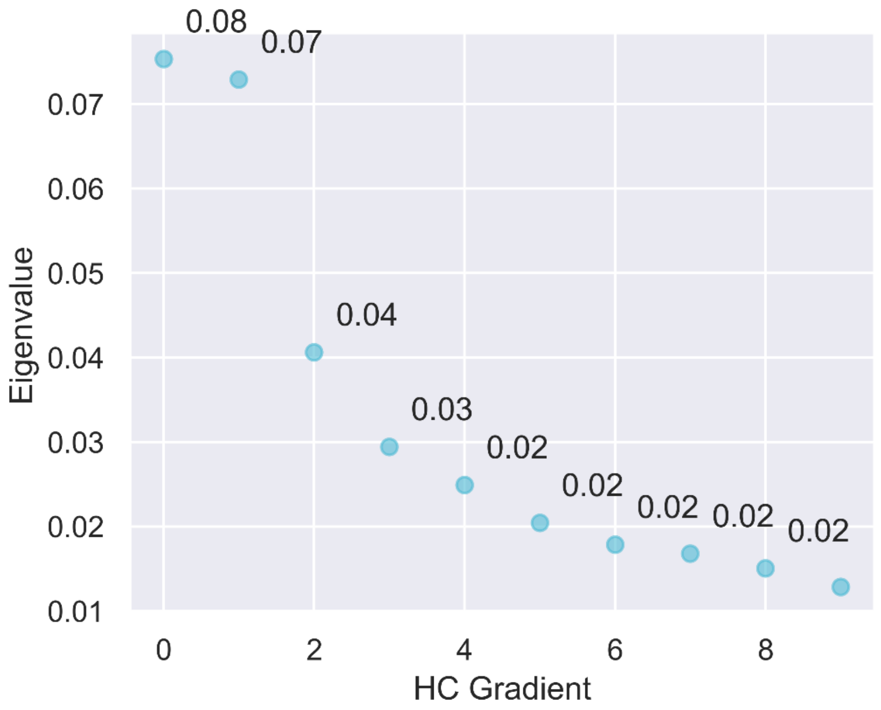


**Fig. S14.** Scree plot for the ten gradients out from diffusion mapping in HC.

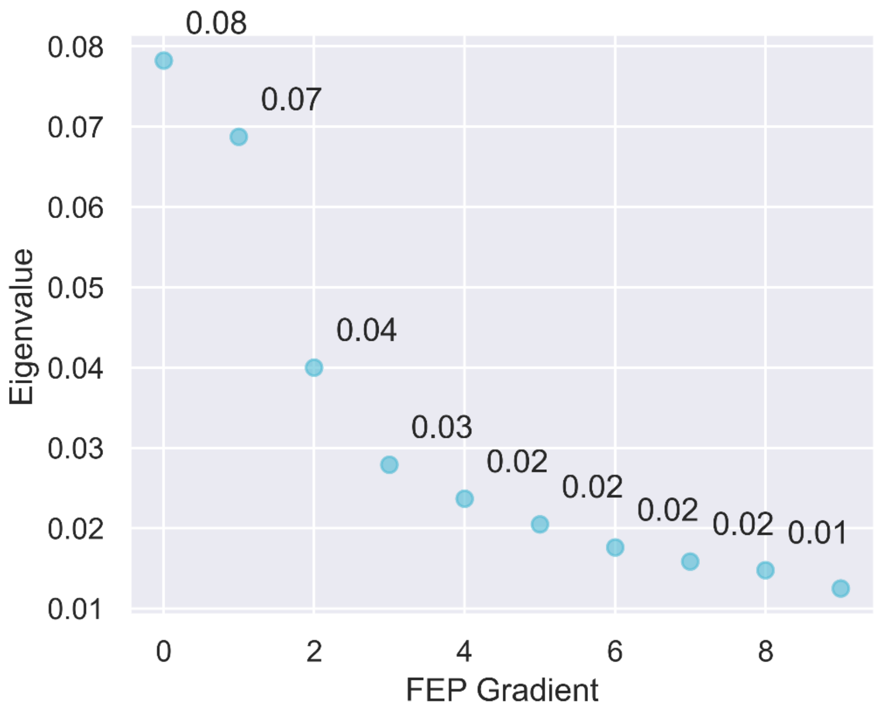


**Fig. S15.** Scree plot for the ten gradients out from diffusion mapping in FEP.

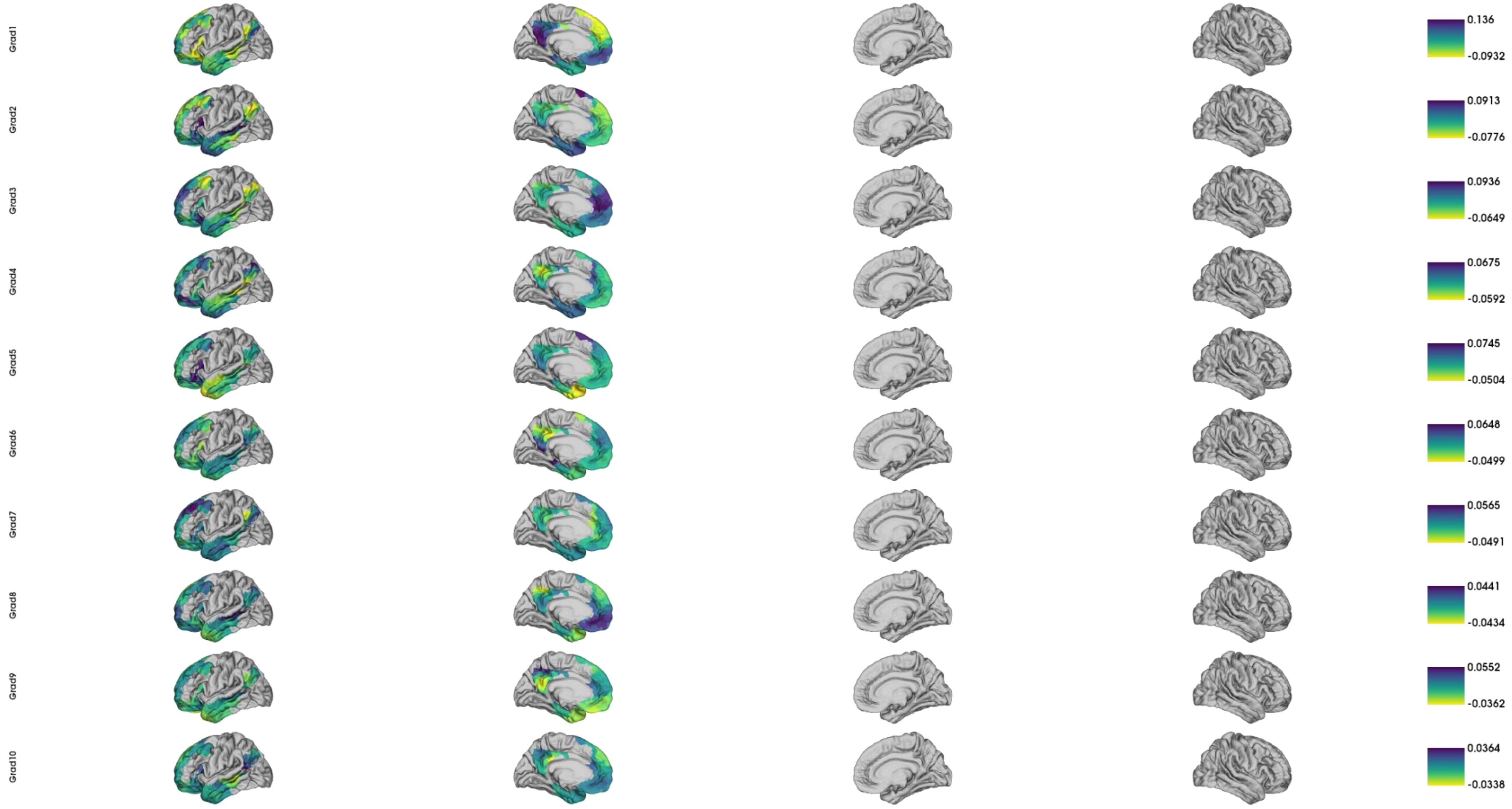


**Fig. S16.** Group templates of semantic network gradients from the principal gradient to the tenth gradient in HC.

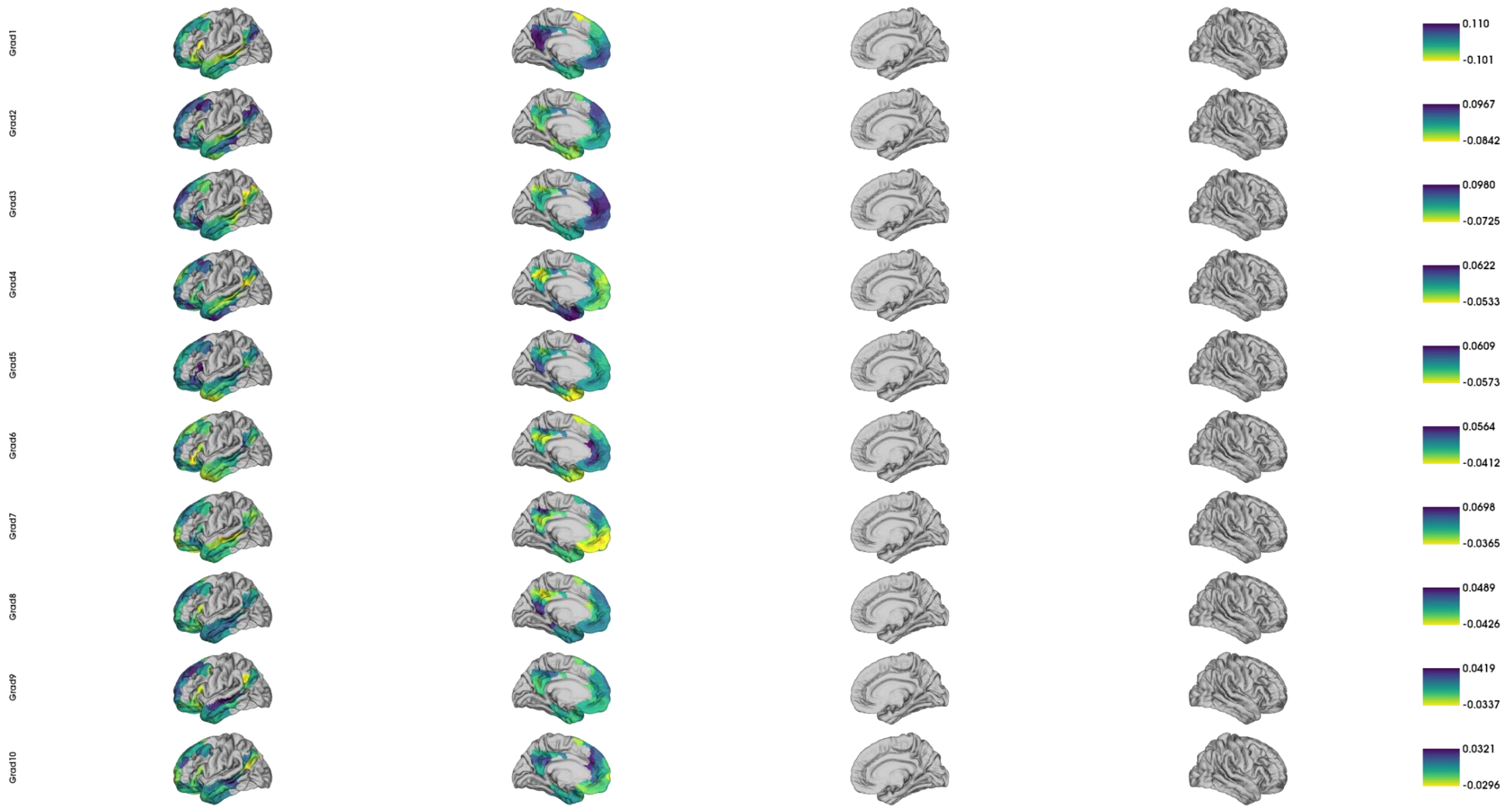


**Fig. S17.** Group templates of semantic network gradients from the principal gradient to the tenth gradient in FEP.

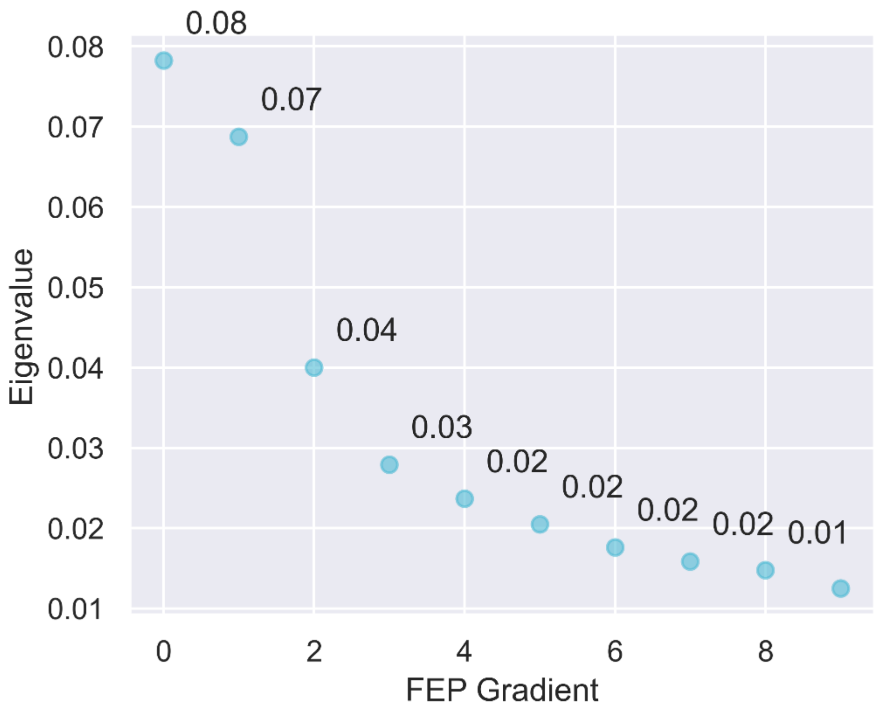


**Fig. S18.** Scree plot for the ten semantic network gradients in HC.

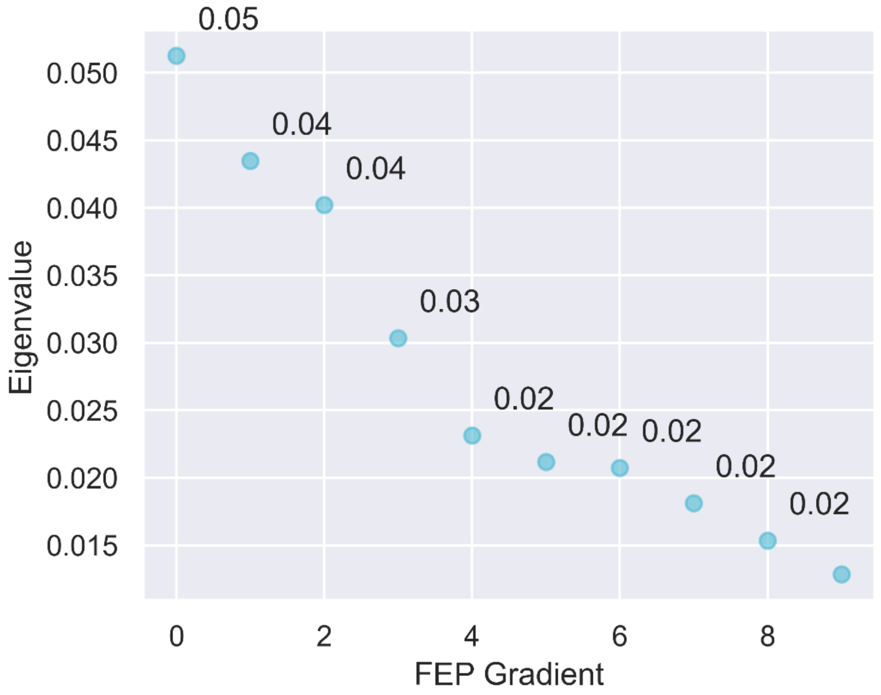


**Fig. S19.** Scree plot for the ten semantic network gradients in FEP.

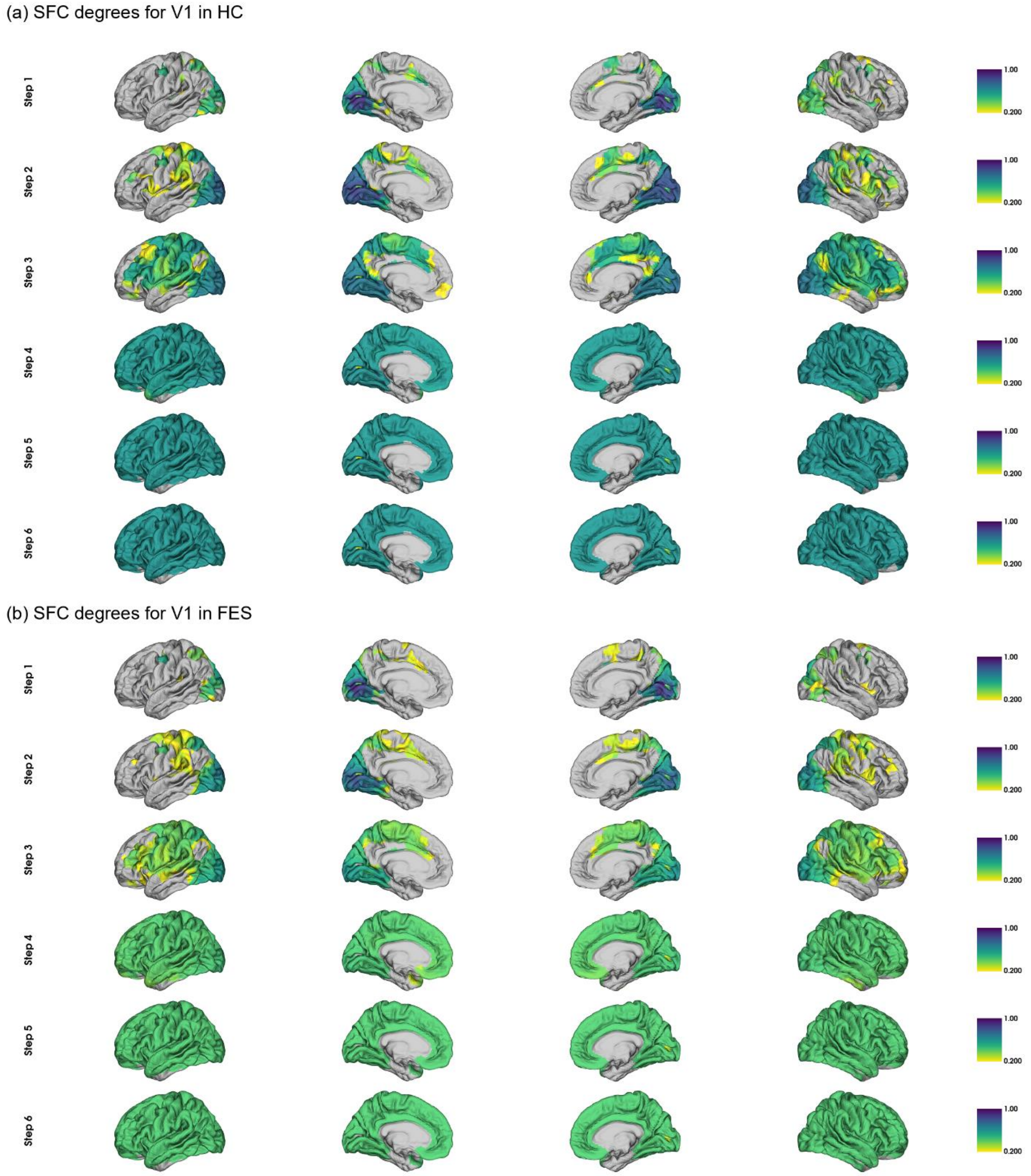


**Fig. S20.** Group-level SFC temples for V1 in (a) HC and (b) FEP in the first six steps (from top to down). To better view the growth of SFC degrees with steps, we only plot SFC degrees larger than 0.2.

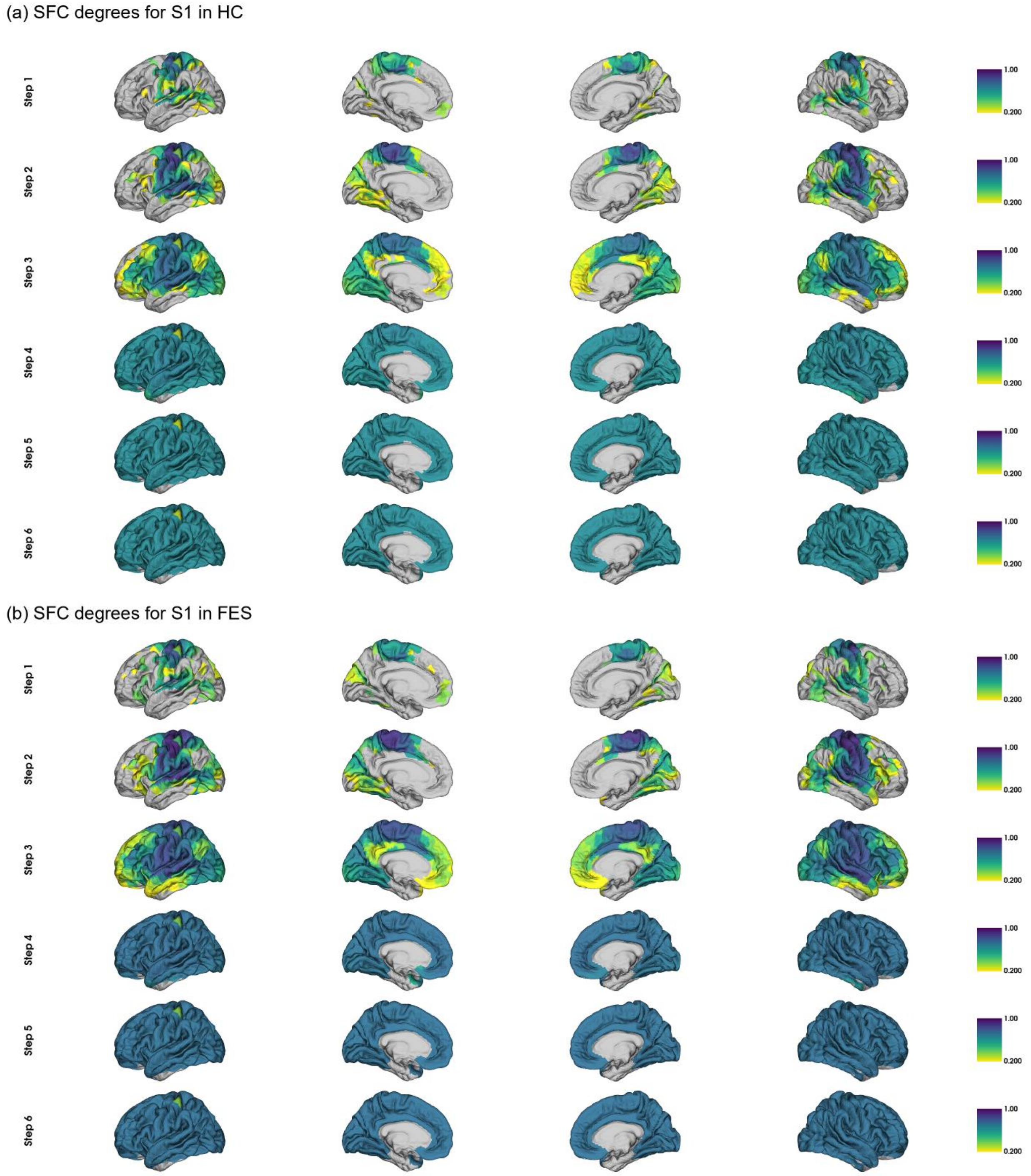


**Fig. S21.** Group-level SFC temples for S1 in (a) HC and (b) FEP in the first six steps (from top to down). To better view the growth of SFC degrees with steps, we only plot SFC degrees larger than 0.2.

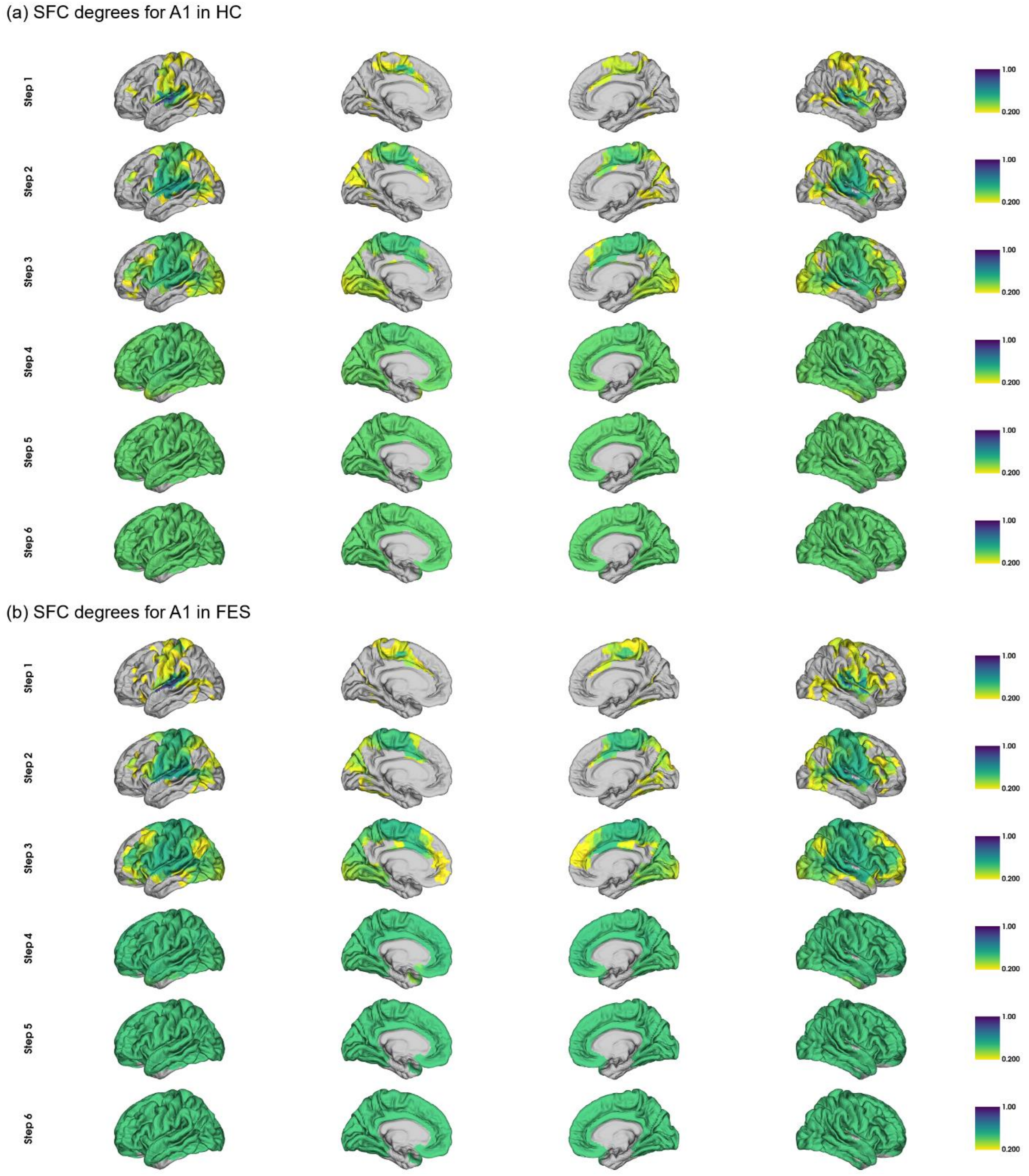


**Fig. S22.** Group-level SFC temples for A1 in (a) HC and (b) FEP in the first six steps (from top to down). To better view the growth of SFC degrees with steps, we only plot SFC degrees larger than 0.2.

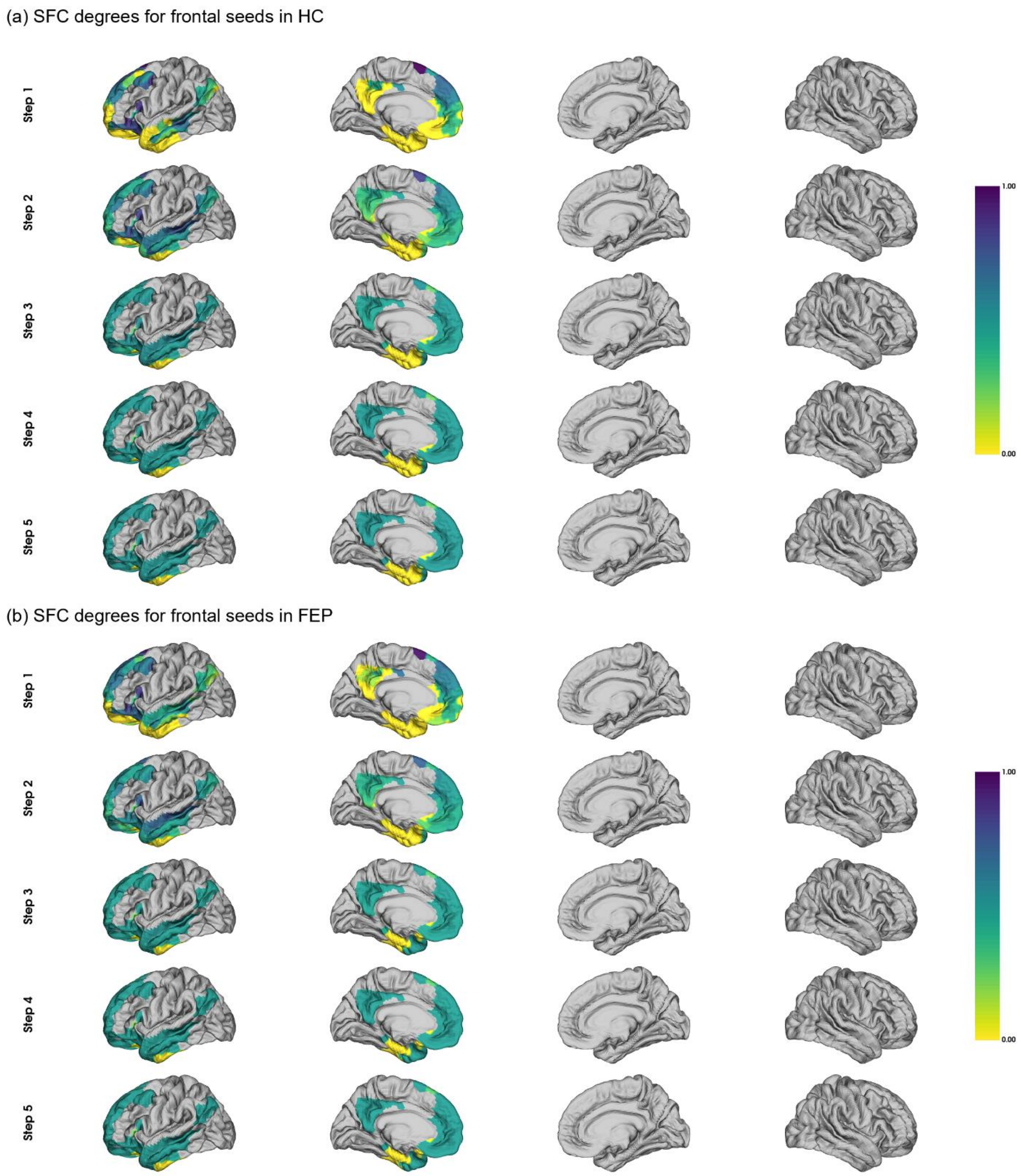


**Fig. S23.** Group-level SFC temples for frontal heteromodal association seeds in (a) HC and (b) FEP in the first five steps (from top to down).

(a) SFC degrees for parietal seeds in HC

(b) SFC degrees for parietal seeds in FEP

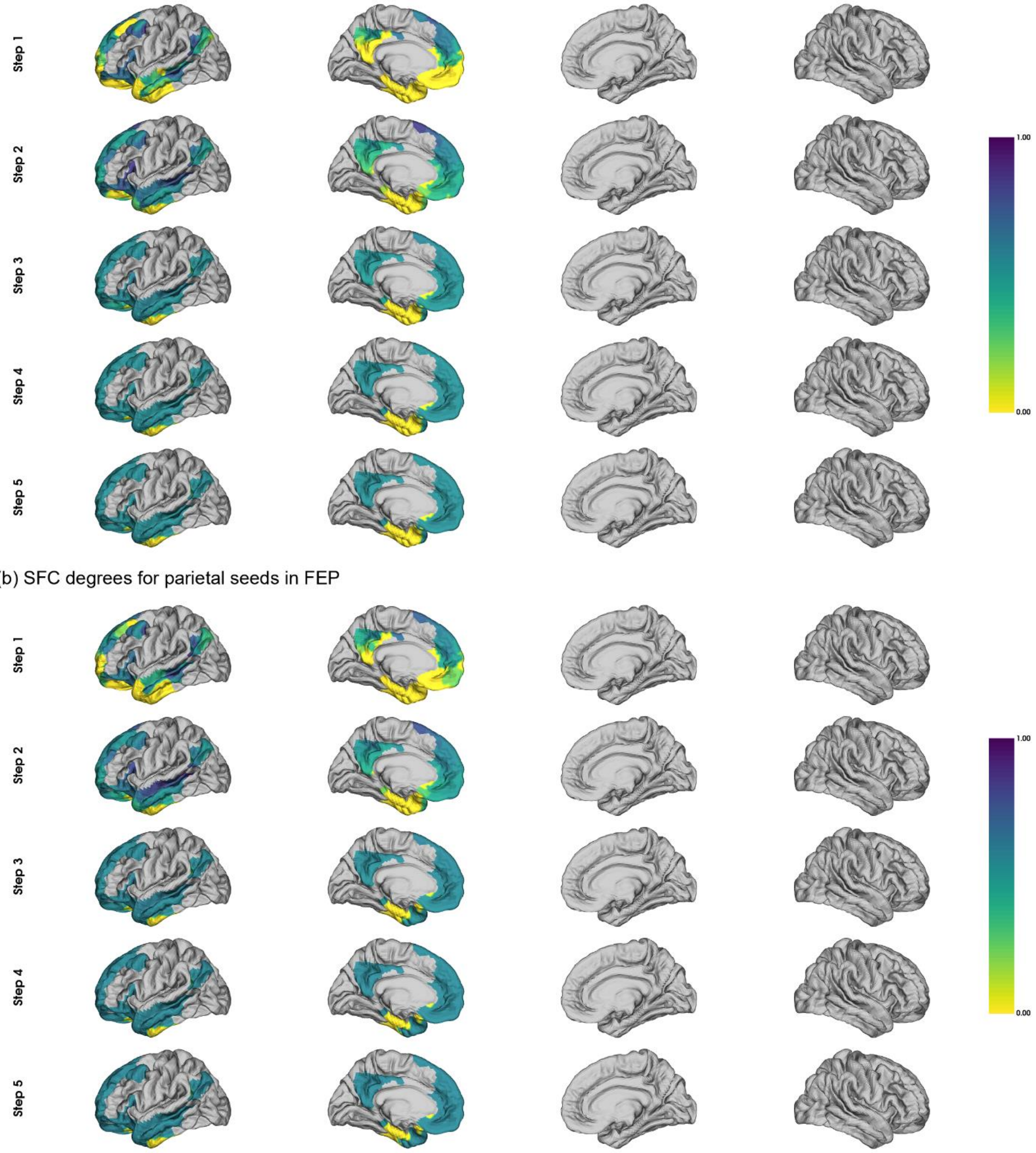


**Fig. S24.** Group-level SFC temples for parietal heteromodal association seeds in (a) HC and (b) FEP in the first six steps (from top to down).

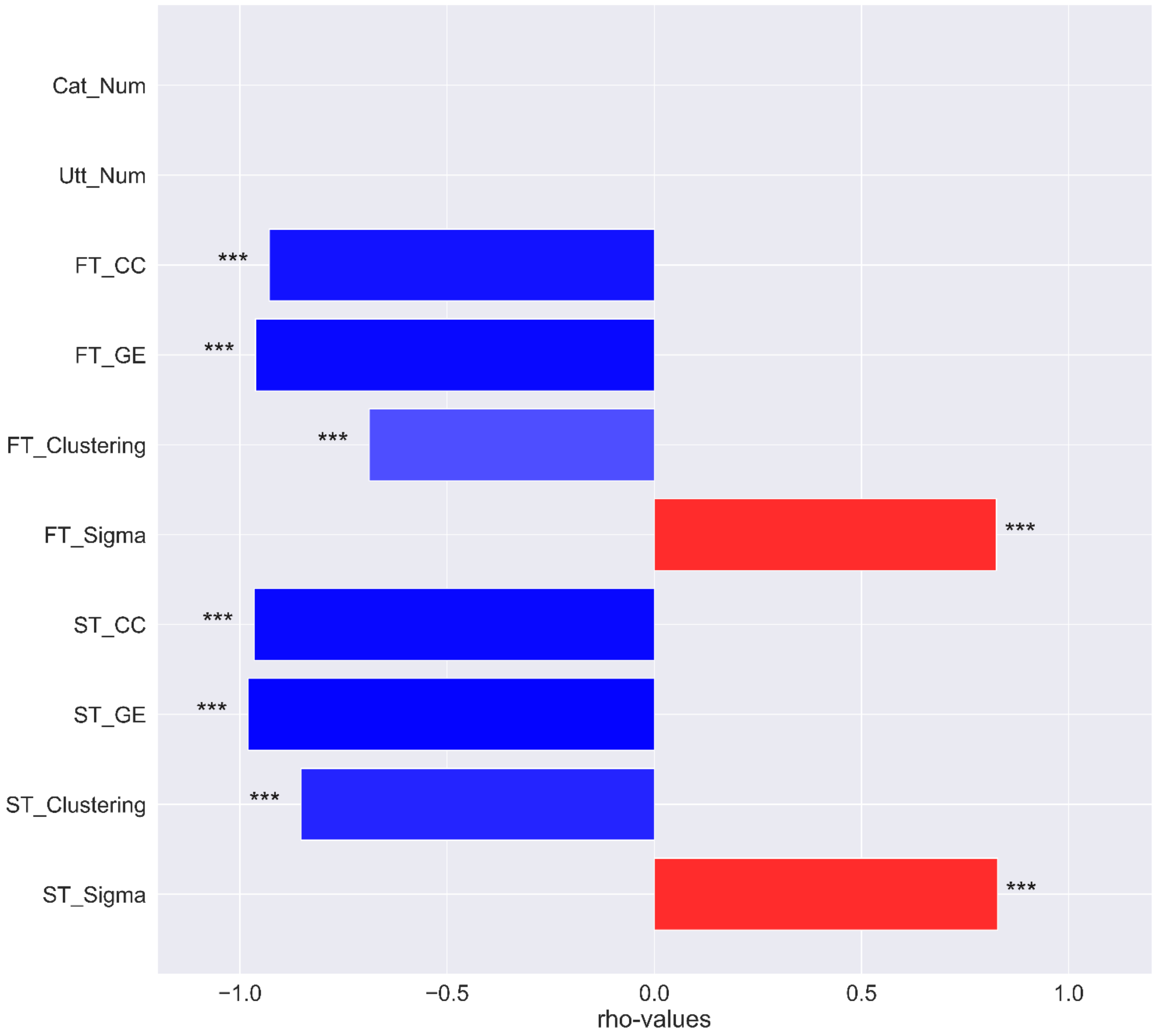


**Fig. S25.** Correlations between semantic measures with the number of lexical categories (FT-measures) or utterances (ST-measures). Red means positive correlations while blue means negative correlations. Significance levels: *** 0.001, ** 0.01, * 0.05.

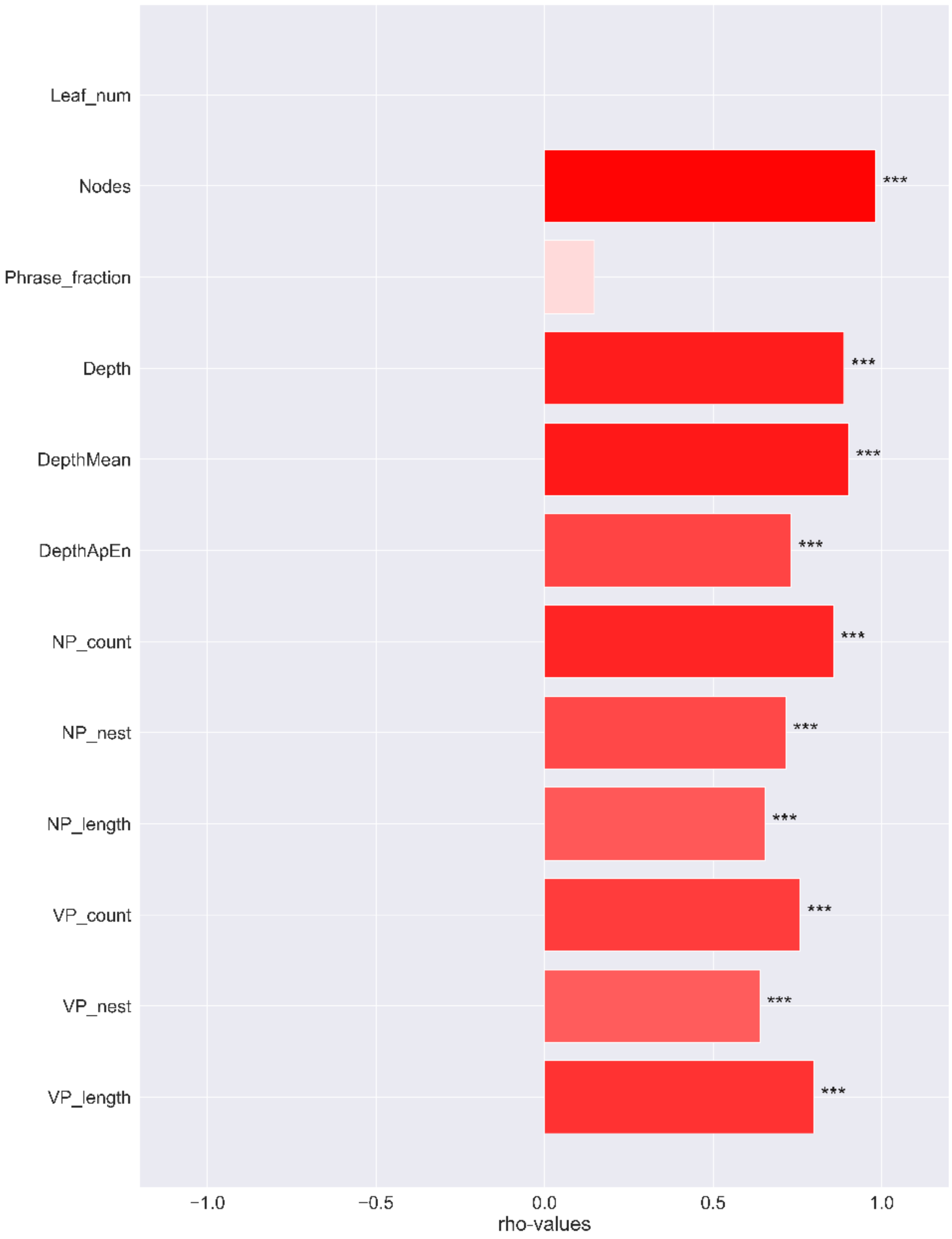


**Fig. S26.** Correlations between syntactic measures with the number of words. Red means positive correlations while blue means negative correlations. Significance levels: *** 0.001, ** 0.01, * 0.05.

**Table S1.** MNI coordinates and corresponding names of SFC ROIs.

| Name | Coordinates (x, y, z) | Name in Schafer's parcellation |
|---|---|---|
| LH V1 | -14, -78, 8 | 7Networks_LH_Vis_47 |
| RH V1 | 10, -78, 8 | 7Networks_RH_Vis_44 |
| LH A1 | -54, -14, 8 | 7Networks_LH_SomMot_5 |
| RH A1 | 58, -14, 8 | 7Networks_RH_SomMot_9 |
| LH S1 | -42, -29, 65 | 7Networks_LH_SomMot_51 |
| RH S1 | 38, -29, 65 | 7Networks_RH_SomMot_78 |

**Table S2.** MNI coordinate and BA names of ROIs with lowest and highest five gradient values in HC and FEP.

| ROI Name | R | A | S | Cortex | BA |
|---|---|---|---|---|---|
| Default_Par_7 | -62 | -54 | 26 | Heteromodal association | AG 39 |
| Default_Par_13 | -60 | -50 | 34 | Heteromodal association | AG 39 |
| Default_PFC_46 | -6 | 24 | 60 | Heteromodal association | FrontEyeFields (8),<br>near dlPFC(dorsal)(9) |
| Default_PFC_17 | -52 | 28 | 2 | Heteromodal association | TriIFG 45 |

Note: AG: angular gyrus; MTG: middle temporal gyrus; TriIFG: triangular part of the inferior frontal gyrus (IFG); PCC: posterior cingulate cortex. The MNI coordinates come from the RAS coordinates as the parcellation is in MNI152 standard space.

**Table S3.** ROIs with the lowest and highest five gradient values in HC and FEP.

| Group | Gradient place | ROI Name | R | A | S | BA | Cortex |
|---|---|---|---|---|---|---|---|
| HC | Lowest 5 | Default_Par_7 | -62 | -54 | 26 | AG 39 | HAC |
| | | Default_Par_13 | -60 | -50 | 34 | AG 39 | HAC |
| | | Default_Temp_14 | -58 | -32 | -4 | MTG 21 | HAC |
| | | Default_PFC_17 | -52 | 28 | 2 | TriIFG 45 | HAC |
| | | Default_PFC_46 | -6 | 24 | 60 | FrontEye 8 (SFG) | HAC |
| | Highest 5 | Default_pCunPCC_2 | -8 | -54 | 8 | AntPFC 10 | MLR |
| | | Default_pCunPCC_4 | -10 | -58 | 16 | VentralPCC 23 | MLR |
| | | Default_pCunPCC_5 | -4 | -54 | 18 | VentralPCC 23 | MLR |
| | | Default_pCunPCC_10 | -4 | -60 | 22 | DorsalPCC 31 | MLR |
| | | Default_pCunPCC_13 | -6 | -66 | 26 | DorsalPCC 31 | MLR |
| FEP | Lowest 5 | Default_Temp_19 | -54 | -42 | 2 | MTG 21 | HAC |
| | | Default_Temp_21 | -54 | -34 | 4 | MTG 21 | HAC |
| | | Default_Temp_22 | -50 | -44 | 8 | MTG 21 | HAC |
| | | Default_PFC_15 | -44 | 28 | -2 | ParsOrbitalis 47 | HAC |
| | | Default_PFC_21 | -54 | 18 | 12 | OpercIFG 44 | HAC |
| | Highest 5 | Default_pCunPCC_4 | -10 | -58 | 16 | VentralPCC 23 | MLR |
| | | Default_pCunPCC_5 | -4 | -54 | 18 | VentralPCC 23 | MLR |
| | | Default_pCunPCC_10 | -4 | -60 | 22 | DorsalPCC 31 | MLR |
| | | Default_pCunPCC_15 | -6 | -58 | 30 | DorsalPCC 31 | MLR |
| | | Default_pCunPCC_18 | -4 | -64 | 32 | DorsalPCC 31 | MLR |

Note: AG: angular gyrus; MTG: middle temporal gyrus; TriIFG: triangular part of the inferior frontal gyrus (IFG); SFG: superior frontal gyrus; AntPFC: anterior prefrontal cortex; PCC: posterior cingulate cortex; ParsOrbitalis: orbital part of IFG; OpercIFG: opercularis part of IFG. HAC: heteromodal association cortex, MLR: medial limbic regions.

**Table S4.** Names, RAS coordinates, cortices, and BA areas of ROIs in the semantic network.

| ROI Name | R | A | S | Cortex | BA (Near) | Note |
|---|---|---|---|---|---|---|
| Default_Par_13 | -60 | -50 | 34 | HAC | AngGyrus (39) | |
| Default_Par_7 | -62 | -54 | 26 | HAC | AngGyrus (39) | |
| Default_PFC_46 | -6 | 24 | 60 | HAC | dlPFC(dorsal) (9) | FrontEyeFields (8), changed to dlPFC(dorsal) (9) |
| Default_PFC_17 | -52 | 28 | 2 | HAC | Broca-Triang (45) | |
| Default_Temp_14 | -58 | -32 | -4 | HAC | MedTempGyrus (21) | |
| Default_Par_11 | -54 | -58 | 34 | HAC | AngGyrus (39) | |
| Default_PFC_8 | -46 | 30 | -10 | HAC | ParsOrbitalis (47) | |
| Default_Temp_10 | -64 | -24 | -8 | HAC | MedTempGyrus (21) | |
| Default_PFC_4 | -38 | 24 | -14 | HAC | ParsOrbitalis (47) | |
| Default_PFC_39 | -2 | 34 | 48 | HAC | dlPFC(dorsal) (9) | FrontEyeFields (8), changed to dlPFC(dorsal) (9) |
| Default_PFC_33 | -4 | 48 | 38 | HAC | dlPFC(dorsal) (9) | |
| Default_PFC_40 | -8 | 42 | 52 | HAC | dlPFC(dorsal) (9) | FrontEyeFields (8), changed to dlPFC(dorsal) (9) |
| Default_PFC_15 | -44 | 28 | -2 | HAC | ParsOrbitalis (47) | |
| Default_Par_6 | -52 | -54 | 26 | HAC | AngGyrus (39) | |
| Default_PFC_21 | -54 | 18 | 12 | HAC | Broca-Operc (44) | |
| Default_Temp_13 | -52 | -22 | -8 | HAC | MedTempGyrus (21) | |
| Default_PFC_11 | -50 | 40 | -6 | HAC | ParsOrbitalis (47) | |
| Default_Temp_16 | -62 | -42 | 0 | HAC | MedTempGyrus (21) | |
| Default_Par_16 | -54 | -60 | 42 | HAC | AngGyrus (39) | Outside defined Bas, changed to AngGyrus (39) |
| Default_PFC_27 | -6 | 50 | 22 | HAC | dlPFC(dorsal) (9) | |
| Default_Temp_17 | -48 | -34 | -2 | HAC | MedTempGyrus (21) | |
| Default_PFC_47 | -14 | 28 | 58 | HAC | dlPFC(dorsal) (9) | FrontEyeFields (8), changed to dlPFC(dorsal) (9) |
| Default_Par_10 | -46 | -58 | 34 | HAC | AngGyrus (39) | |
| Default_Par_18 | -52 | -50 | 42 | HAC | AngGyrus (39) | |
| Default_PFC_49 | -4 | 12 | 64 | HAC | dlPFC(dorsal) (9) | PreMot+SuppMot (6), changed to dlPFC(dorsal) (9) |
| Default_Temp_19 | -54 | -42 | 2 | HAC | MedTempGyrus (21) | |
| Default_Temp_12 | -66 | -36 | -6 | HAC | MedTempGyrus (21) | |
| Default_PFC_32 | -4 | 36 | 34 | HAC | dlPFC(dorsal) (9) | FrontEyeFields (8), changed to dlPFC(dorsal) (9) |
| Default_PFC_42 | -42 | 6 | 48 | HAC | dlPFC(dorsal) (9) | PreMot+SuppMot (6), changed to dlPFC(dorsal) (9) |
| Default_PFC_26 | -24 | 58 | 20 | HAC | AntPFC (10) | |

| | | | | | | |
|---|---|---|---|---|---|---|
| Default_PFC_37 | -38 | 24 | 46 | HAC | dlPFC(dorsal) (9) | FrontEyeFields (8), changed to dlPFC(dorsal) (9) |
| Default_PFC_31 | -22 | 50 | 30 | HAC | AntPFC (10) | |
| Default_PFC_30 | -6 | 62 | 28 | HAC | AntPFC (10) | |
| Default_Par_3 | -60 | -52 | 18 | HAC | AngGyrus (39) | |
| Default_Temp_20 | -60 | -50 | 6 | HAC | MedTempGyrus (21) | |
| Default_PFC_50 | -16 | 14 | 66 | HAC | dlPFC(dorsal) (9) | PreMot+SuppMot (6), changed to dlPFC(dorsal) (9) |
| Default_PFC_6 | -38 | 50 | -14 | HAC | AntPFC (10) | |
| Default_Temp_21 | -54 | -34 | 4 | HAC | MedTempGyrus (21) | |
| Default_pCunPCC_21 | -2 | -14 | 36 | MLR | VentPostCing (23) | |
| Default_PFC_18 | -44 | 20 | 6 | HAC | Broca-Triang (45) | |
| Default_Temp_22 | -50 | -44 | 8 | HAC | MedTempGyrus (21) | |
| Default_PFC_14 | -28 | 58 | -2 | HAC | AntPFC (10) | |
| Default_PFC_36 | -46 | 14 | 40 | HAC | dlPFC(dorsal) (9) | FrontEyeFields (8), changed to dlPFC(dorsal) (9) |
| Default_Temp_1 | -46 | 8 | -36 | HAC | Temporalpole (38) | |
| Default_Par_5 | -44 | -56 | 26 | HAC | AngGyrus (39) | |
| Default_Par_1 | -48 | -58 | 18 | HAC | AngGyrus (39) | |
| Default_Temp_8 | -62 | -34 | -14 | HAC | MedTempGyrus (21) | |
| Default_Temp_5 | -48 | 14 | -24 | HAC | Temporalpole (38) | |
| Default_PFC_29 | -16 | 60 | 26 | HAC | AntPFC (10) | |
| Default_Temp_18 | -58 | -24 | 0 | HAC | SupTempGyrus (22) | |
| Default_Temp_2 | -52 | -6 | -32 | HAC | InfTempGyrus (20) | |
| Default_Par_15 | -46 | -62 | 42 | HAC | AngGyrus (39) | |
| Limbic_OFC_14 | -22 | 62 | -10 | MLR | OrbFrontal (11) | AntPFC (10), changed to OrbFrontal (11) |
| Default_PFC_1 | -28 | 16 | -16 | HAC | ParsOrbitalis (47) | Insula (13), changed to ParsOrbitalis (47) |
| Default_Temp_7 | -52 | 6 | -16 | HAC | Temporalpole (38) | |
| Default_Par_2 | -46 | -50 | 20 | HAC | AngGyrus (39) | |
| Limbic_OFC_13 | -10 | 62 | -16 | MLR | OrbFrontal (11) | |
| Default_PFC_34 | -12 | 52 | 40 | HAC | dlPFC(dorsal) (9) | |
| Default_PFC_28 | -8 | 36 | 20 | HAC | DorsalACC (32) | |
| Default_Temp_15 | -58 | -14 | -4 | HAC | SupTempGyrus (22) | |
| Default_Par_19 | -44 | -64 | 52 | HAC | AngGyrus (39) | |
| Default_Temp_4 | -54 | 4 | -26 | HAC | Temporalpole (38) | |
| Default_Temp_3 | -60 | -22 | -26 | HAC | InfTempGyrus (20) | |
| Default_Temp_9 | -60 | -10 | -14 | HAC | MedTempGyrus (21) | |

| | | | | | | |
|---|---|---|---|---|---|---|
| Default_PFC_20 | -6 | 44 | 6 | HAC | DorsalACC (32) | |
| Default_PFC_19 | -6 | 60 | 8 | HAC | AntPFC (10) | |
| Limbic_OFC_8 | -18 | 40 | -20 | MLR | OrbFrontal (11) | |
| Default_PFC_35 | -20 | 44 | 42 | HAC | dlPFC(dorsal) (9) | |
| Limbic_OFC_10 | -8 | 50 | -22 | MLR | OrbFrontal (11) | |
| Default_PFC_43 | -38 | 14 | 56 | HAC | dlPFC(dorsal) (9) | PreMot+SuppMot (6), changed to dlPFC(dorsal) (9) |
| Default_PFC_25 | -2 | 30 | 18 | HAC | AntPFC (10) | VentAntCing (24), changed to AntPFC (10) |
| Limbic_OFC_5 | -28 | 16 | -24 | MLR | OrbFrontal (11) | ParsOrbitalis (47), changed to OrbFrontal (11) |
| Default_PFC_2 | -34 | 12 | -12 | HAC | ParsOrbitalis (47) | Insula (13), changed to ParsOrbitalis (47) |
| Default_Temp_11 | -54 | -4 | -10 | HAC | SupTempGyrus (22) | |
| Limbic_TempPole_15 | -44 | 6 | -16 | MLR | Temporalpole (38) | |
| Limbic_TempPole_8 | -36 | 18 | -34 | MLR | Temporalpole (38) | |
| Limbic_TempPole_5 | -50 | -14 | -36 | MLR | InfTempGyrus (20) | |
| Limbic_TempPole_11 | -54 | -32 | -28 | MLR | InfTempGyrus (20) | |
| Default_pCunPCC_22 | -2 | -26 | 38 | MLR | VentPostCing (23) | |
| Limbic_TempPole_9 | -26 | 8 | -30 | MLR | Temporalpole (38) | |
| Limbic_TempPole_4 | -28 | -8 | -34 | MLR | Parahipp (36) | |
| Limbic_OFC_1 | -20 | 12 | -22 | MLR | OrbFrontal (11) | |
| Limbic_TempPole_14 | -38 | 8 | -24 | MLR | Temporalpole (38) | |
| Limbic_TempPole_3 | -38 | -12 | -38 | MLR | InfTempGyrus (20) | |
| Limbic_TempPole_1 | -28 | 2 | -44 | MLR | Temporalpole (38) | |
| Limbic_TempPole_2 | -42 | -2 | -44 | MLR | Temporalpole (38) | Outside defined Bas, changed to Temporalpole (38) |
| Default_Par_17 | -38 | -72 | 48 | HAC | AngGyrus (39) | |
| Limbic_TempPole_6 | -18 | -2 | -30 | MLR | Parahipp (36) | |
| Default_Par_9 | -48 | -64 | 32 | HAC | AngGyrus (39) | |
| Default_Temp_6 | -62 | -16 | -20 | HAC | MedTempGyrus (21) | |
| Limbic_TempPole_10 | -44 | -30 | -22 | MLR | InfTempGyrus (20) | |
| Limbic_OFC_3 | -14 | 24 | -24 | MLR | OrbFrontal (11) | |
| Default_PFC_48 | -22 | 22 | 58 | HAC | dlPFC(dorsal) (9) | PreMot+SuppMot (6), changed to dlPFC(dorsal) (9) |
| Default_PFC_24 | -8 | 68 | 16 | HAC | AntPFC (10) | |
| Limbic_TempPole_7 | -40 | -18 | -28 | MLR | InfTempGyrus (20) | |
| Default_Par_14 | -46 | -70 | 40 | HAC | AngGyrus (39) | |
| Default_PFC_44 | -30 | 16 | 56 | HAC | dlPFC(dorsal) (9) | PreMot+SuppMot (6), changed to dlPFC(dorsal) (9) |
| Default_PFC_41 | -20 | 34 | 50 | HAC | dlPFC(dorsal) (9) | FrontEyeFields (8), changed to dlPFC(dorsal) (9) |

| | | | | | | |
|---|---|---|---|---|---|---|
| Default_PFC_23 | -2 | 36 | 8 | HAC | AntPFC (10) | VentAntCing (24), changed to AntPFC (10) |
| Default_PFC_9 | -4 | 24 | -6 | HAC | AntPFC (10) | VentAntCing (24), changed to AntPFC (10) |
| Default_PFC_12 | -16 | 66 | -4 | HAC | AntPFC (10) | |
| Default_PFC_22 | -20 | 66 | 10 | HAC | AntPFC (10) | |
| Limbic_OFC_7 | -24 | 28 | -18 | MLR | OrbFrontal (11) | |
| Default_PFC_45 | -22 | 18 | 46 | HAC | dlPFC(dorsal) (9) | FrontEyeFields (8), changed to dlPFC(dorsal) (9) |
| Default_pCunPCC_25 | -4 | -32 | 42 | MLR | DorsalPCC (31) | |
| Limbic_OFC_2 | -12 | 22 | -18 | MLR | OrbFrontal (11) | |
| Limbic_TempPole_12 | -20 | -14 | -30 | MLR | Parahipp (36) | |
| Default_pCunPCC_31 | -6 | -48 | 44 | MLR | DorsalPCC (31) | |
| Limbic_OFC_4 | -10 | 34 | -22 | MLR | OrbFrontal (11) | |
| Default_PFC_38 | -26 | 28 | 44 | HAC | dlPFC(dorsal) (9) | FrontEyeFields (8), changed to dlPFC(dorsal) (9) |
| Limbic_OFC_12 | -4 | 16 | -10 | MLR | OrbFrontal (11) | Subgenual (25), changed to OrbFrontal (11) |
| Default_pCunPCC_27 | -4 | -40 | 42 | MLR | DorsalPCC (31) | |
| Default_PFC_3 | -36 | 36 | -12 | HAC | ParsOrbitalis (47) | |
| Default_PHC_2 | -20 | -24 | -22 | MLR | Parahipp (36) | |
| Default_PFC_13 | -4 | 36 | -4 | HAC | DorsalACC (32) | |
| Default_Par_4 | -44 | -64 | 24 | HAC | AngGyrus (39) | |
| Default_pCunPCC_29 | -6 | -54 | 42 | MLR | DorsalPCC (31) | |
| Limbic_TempPole_13 | -32 | -24 | -26 | MLR | Parahipp (36) | |
| Default_pCunPCC_11 | -4 | -42 | 26 | MLR | VentPostCing (23) | |
| Limbic_OFC_6 | -8 | 20 | -22 | MLR | OrbFrontal (11) | |
| Default_pCunPCC_16 | -4 | -40 | 30 | MLR | VentPostCing (23) | |
| Default_pCunPCC_26 | -12 | -48 | 38 | MLR | DorsalPCC (31) | |
| Default_pCunPCC_32 | -6 | -58 | 48 | MLR | DorsalPCC (31) | VisMotor (7), chagned to DorsalPCC (31) |
| Default_pCunPCC_30 | -4 | -68 | 44 | MLR | DorsalPCC (31) | VisMotor (7), chagned to DorsalPCC (31) |
| Limbic_OFC_11 | -2 | 20 | -18 | MLR | OrbFrontal (11) | |
| Default_pCunPCC_20 | -10 | -42 | 34 | MLR | VentPostCing (23) | |
| Default_pCunPCC_17 | -12 | -50 | 30 | MLR | VentPostCing (23) | |
| Default_PFC_16 | -8 | 68 | 2 | HAC | AntPFC (10) | |
| Default_pCunPCC_6 | -6 | -44 | 20 | MLR | VentPostCing (23) | |
| Default_pCunPCC_1 | -6 | -46 | 8 | MLR | AgrRetrolimb (30) | |
| Default_PFC_10 | -8 | 46 | -4 | HAC | DorsalACC (32) | |
| Default_pCunPCC_14 | -6 | -46 | 30 | MLR | VentPostCing (23) | |

| | | | | | | |
|---|---|---|---|---|---|---|
| Limbic_OFC_9 | -4 | 38 | -22 | MLR | OrbFrontal (11) | |
| Default_Par_12 | -38 | -78 | 40 | HAC | AngGyrus (39) | |
| Default_PHC_1 | -30 | -32 | -18 | MLR | Fusiform (37) | |
| Default_pCunPCC_3 | -6 | -46 | 14 | MLR | AgrRetrolimb (30) | |
| Default_pCunPCC_9 | -16 | -68 | 26 | MLR | DorsalPCC (31) | VisualAssoc (19), changed to DorsalPCC (31) |
| Default_PFC_5 | -6 | 32 | -12 | HAC | DorsalACC (32) | |
| Default_pCunPCC_19 | -4 | -34 | 36 | MLR | VentPostCing (23) | |
| Default_pCunPCC_23 | -8 | -54 | 34 | MLR | DorsalPCC (31) | |
| Default_PHC_3 | -20 | -36 | -14 | MLR | Fusiform (37) | |
| Default_pCunPCC_28 | -6 | -62 | 38 | MLR | DorsalPCC (31) | |
| Default_Par_8 | -48 | -70 | 30 | HAC | AngGyrus (39) | |
| Default_PFC_7 | -4 | 54 | -10 | HAC | AntPFC (10) | |
| Default_pCunPCC_24 | -2 | -70 | 36 | MLR | DorsalPCC (31) | VisMotor (7), chagned to DorsalPCC (31) |
| Default_pCunPCC_8 | -2 | -48 | 24 | MLR | VentPostCing (23) | |
| Default_pCunPCC_7 | -16 | -64 | 22 | MLR | DorsalPCC (31) | SecVisual (18), changed to DorsalPCC (31) |
| Default_pCunPCC_18 | -4 | -64 | 32 | MLR | DorsalPCC (31) | |
| Default_pCunPCC_12 | -8 | -52 | 26 | MLR | VentPostCing (23) | |
| Default_pCunPCC_15 | -6 | -58 | 30 | MLR | DorsalPCC (31) | |
| Default_pCunPCC_2 | -8 | -54 | 8 | MLR | VentPostCing (23) | |
| Default_pCunPCC_5 | -4 | -54 | 18 | MLR | VentPostCing (23) | |
| Default_pCunPCC_4 | -10 | -58 | 16 | MLR | VentPostCing (23) | |
| Default_pCunPCC_13 | -6 | -66 | 26 | MLR | DorsalPCC (31) | |
| Default_pCunPCC_10 | -4 | -60 | 22 | MLR | DorsalPCC (31) | |

HAC: heteromodal heteromodal association cortex, MLR: medial limbic regions.
